# PANDA: Phased Analysis of DNA Amplicons for Read-Level Methylation Profiling and Heterogeneity Assessment

**DOI:** 10.64898/2026.04.01.715790

**Authors:** Azusa Kubota, Hisato Kobayashi, Atsushi Tajima

## Abstract

Targeted bisulfite amplicon sequencing is widely used to examine methylation at defined genomic loci, but site-wise methylation estimates can obscure phased patterns among individual reads. We developed PANDA (Phased Analysis for DNA Amplicons), an open-source platform for read-level analysis of targeted bisulfite amplicon NGS, with additional support for Sanger-derived sequences and graphical and command-line interfaces built on a shared computational core (https://github.com/kubo-azu/PANDA). PANDA integrates reference-coordinate methylation calling and visualization, motif-based haplotype filtering, coordinate linkage of unmerged paired reads, and group comparison. It also implements Amplicon PDR, Window Epipolymorphism, and Amplicon qFDRP with retained read-count weighting and explicit eligibility criteria. Synthetic demonstrations illustrated the analysis functions, while controlled benchmarks evaluated read retention, CpG-call accuracy, heterogeneity metrics, and computational scaling against known simulation truth. Matched-mean populations showed that the metrics capture different properties of observed reads, although qFDRP did not distinguish CpG linkage patterns with identical site-wise frequencies and complete coverage. Re-analysis of public chimpanzee and rhesus macaque placental datasets was consistent with the previously reported species-associated methylation contrast and demonstrated coordinate-linked analysis of unmerged paired reads without imputing the unsequenced interval. PANDA provides a reproducible framework for examining phased methylation patterns and within-sample heterogeneity alongside site-wise methylation estimates.

## Introduction

DNA methylation is a fundamental epigenetic modification that contributes to gene regulation, cellular identity, and genome stability (Mattei et al., 2022). Bisulfite sequencing and related enzymatic conversion methods provide quantitative, single-base-resolution measurements of DNA methylation (Meissner et al., 2005; Masser et al., 2013; Adusumalli et al., 2015; Vaisvila et al., 2021). Recent high-throughput and long-read technologies have extended methylation profiling to genome-wide and long-range applications (Flusberg et al., 2010; Fu et al., 2025; Liu and Conesa, 2025; Montano and Timp, 2025). Nevertheless, targeted bisulfite amplicon sequencing (TBAS) remains valuable for cost-efficient, high-depth analysis of predefined loci across large sample series. It supports routine locus-specific profiling and focused validation of findings from genome-wide studies.

Beyond these practical advantages, targeted bisulfite sequencing preserves methylation information at the level of individual reads or clones. Historically, locus-specific methylation was examined by Sanger sequencing of cloned PCR products, enabling direct visualization of CpG methylation patterns in individual clones (Lewin et al., 2004). The subsequent adoption of next-generation sequencing greatly increased throughput and enabled deeper sampling of phased methylation patterns and epiallelic diversity (Smith et al., 2025). Although these approaches operate at different scales, both retain information about coordinated methylation states across CpGs observed within the same sequence. Sanger data comprise a relatively small number of explicitly represented clones, whereas NGS data comprise many reads whose retained abundance can be incorporated into quantitative summaries.

Molecule-resolved analysis is particularly informative because similar mean methylation levels can arise from fundamentally different molecular compositions. For example, an intermediate methylation level may reflect coordinated allele-specific methylation (ASM) at an imprinted locus (Meaburn et al., 2010; Tycko, 2010; Rosenski et al., 2025), a mixture of epigenetically distinct cell populations, or the accumulation of stochastic and partially discordant methylation patterns (Landan et al., 2012; Jenkinson et al., 2017). Such heterogeneity is also biologically relevant beyond imprinting: epigenetic dysregulation and drift increase with aging (Wang et al., 2022; Yücel and Gladyshev, 2026), whereas tumors often contain diverse methylation states associated with clonal evolution and cellular plasticity (Zoghbi and Beaudet, 2016; Laisné et al., 2025). Site-wise averages alone cannot distinguish among these possibilities. Molecule-level methylation patterns and within-sample heterogeneity should therefore be evaluated alongside CpG-wise methylation percentages. Importantly, heterogeneity metrics are descriptive summaries rather than independent evidence of ASM or another specific mechanism; their interpretation requires integration with methylation distributions, phased patterns, sequence-defined haplotypes, replicate structure, and the experimental context.

Existing software addresses different experimental formats and stages of bisulfite-sequencing analysis. For example, QUMA (Kumaki et al., 2008) and BiQ Analyzer (Bock et al., 2005) support alignment, quality control, methylation quantification, and clone-level visualization for conventional bisulfite cloning and Sanger sequencing. BiQ Analyzer HT (Lutsik et al., 2011) extends locus-specific analysis to high-throughput bisulfite sequencing and provides alignment, quality control, read-level methylation visualization, data export, and graphical and command-line operation. Genome-oriented aligners, including Bismark (Krueger and Andrews, 2011), BS-Seeker2 (Guo et al., 2013), and BiSulfite Bolt (Farrell et al., 2021), map bisulfite NGS reads to reference genomes and generate coordinate-level methylation calls. For targeted NGS data, ampliMethProfiler (Scala et al., 2016) extracts epihaplotype profiles from reads analyzed against user-provided amplicon references, whereas MethPat (Wong et al., 2016) visualizes methylation patterns from Bismark-aligned data and epialleleR (Nikolaienko et al., 2023) reports methylation levels, epiallele frequencies, methylation patterns, and allele specificity from aligned BAM files.

Although these tools provide read-level outputs, their workflows are generally organized around a specific input type or analysis stage. This separation can complicate the analysis of sequence-defined haplotypes or paired reads covering different parts of a long amplicon. A common reference-coordinate representation, appropriate treatment of NGS read abundance, and explicit eligibility rules are needed to compare phased patterns and heterogeneity summaries across samples.

To address these gaps, we developed PANDA (Phased Analysis for DNA Amplicons), a platform for phased, read-level analysis of targeted bisulfite amplicons. An overview of the PANDA workflow is shown in Figure 1. PANDA represents dereplicated amplicon-NGS sequence variants as CpG calls in reference coordinates while retaining observed read abundance; Sanger-derived clone sequences can also be processed in this framework. Its Shiny GUI and CLI use the shared PANDAcore engine for read-level visualization and heterogeneity analysis across interactive and scripted workflows. Table 1 compares PANDA’s functions with those of representative tools for targeted bisulfite analysis.

**Figure 1.**
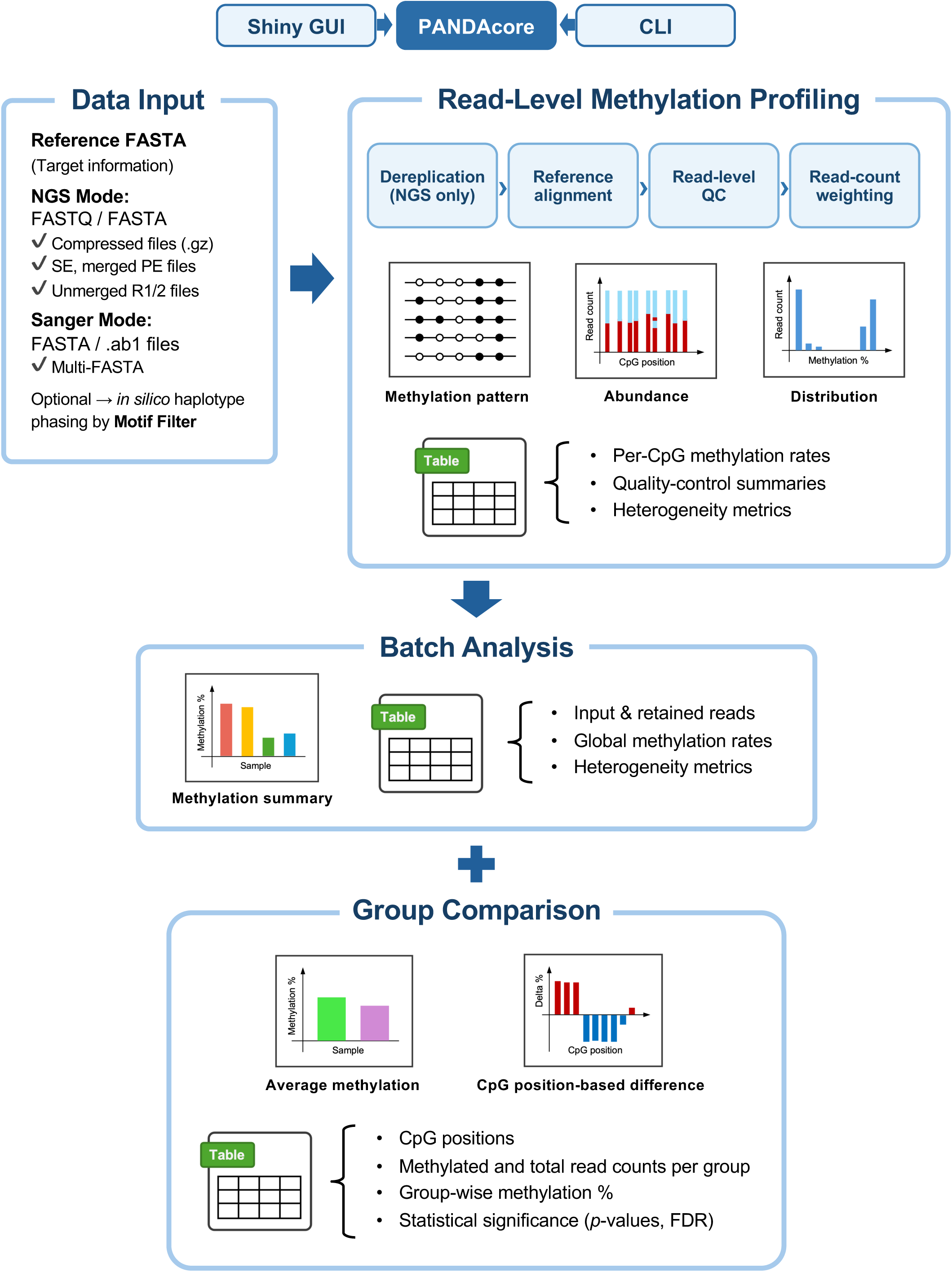
Overview of the PANDA platform and analysis workflow. The Shiny graphical interface and command-line interface use the same PANDAcore functions. PANDA accepts a reference FASTA together with amplicon-NGS reads in FASTA or FASTQ format, or Sanger-derived sequences in AB1 or FASTA format. NGS inputs may consist of single-end, merged, or unmerged paired reads. The workflow includes exact-sequence dereplication of NGS reads, bisulfite-aware reference alignment, read-level quality control, and retention of read counts as weights in downstream summaries. Read-level analysis generates phased methylation patterns, abundance heatmaps, methylation distributions, quality-control summaries, and three heterogeneity metrics: Amplicon PDR, Window Epipolymorphism, and Amplicon qFDRP. Batch analysis summarizes results across multiple samples, whereas group comparison evaluates overall and CpG-level methylation differences. Optional motif filtering supports analysis of sequence-defined haplotypes.

**Table 1.** Representative tools for read-level analysis of targeted bisulfite sequencing data.

| Tool | Input | Read-level output | Pattern abundance | Sequence-defined haplotypes | Heterogeneity analysis | Paired-reads | Interface |
|---|---|---|---|---|---|---|---|
| QUMA | Bisulfite-sequenced clones | Yes | – | – | – | Not applicable | Web / local |
| BiQ Analyzer | Bisulfite-sequenced clones | Yes | – | – | – | Not applicable | Desktop GUI |
| BiQ Analyzer HT | Targeted FASTA / aligned BAM | Yes | Per-read output | – | – | Not specified | GUI / CLI |
| ampliMethProfiler | FASTA + amplicon reference | Yes | Epihaplotype counts | – | Epihaplotype α/β diversity | Merged upstream | CLI |
| MethPat | Bismark output + regions | Yes | Pattern counts | – | – | Merged upstream | CLI / HTML |
| epialleleR | Aligned BAM | Yes | Epiallele frequencies | SNV–methylation association (with VCF) | Epiallele frecuencies | Aligned pairs | R package |
| **PANDA** (this tool) | Reference FASTA +<br>AB1 / FASTA / FASTQ | Yes | Read count-weighted patterns | Motif filtering | Amplicon PDR, Window Epipolymorphism, Amplicon qFDRP | Merged / unmerged paired reads | GUI / CLI |

PDR, proportion of discordant reads; qFDRP, quantitative fraction of discordant read pairs; GUI, graphical user interface; CLI, command-line interface. “Per-read output” denotes individual read records rather than aggregated pattern counts; “–” indicates that the specified native function was not documented.

## Method

### Software implementation and workflow

PANDA is implemented in R, with PANDAcore providing the computational package shared by the Shiny GUI and CLI. The software environment and direct package dependencies used for the analyses reported here are summarized in Table S1. The CLI provides `panda analyze` for analysis, `panda plot` for figure generation, and `panda compare` for predefined two-group comparisons; these commands produce CSV, JSON, and PDF outputs.

### Data input and motif-based haplotype filtering

Analyses begin with a reference amplicon FASTA; users select the target when it contains multiple records. Sanger mode accepts FASTA, multi-FASTA, and AB1-derived sequences. Amplicon-NGS mode accepts FASTA and compressed or uncompressed FASTQ from single-end, pre-merged paired-end, or explicitly paired unmerged R1/R2 inputs.

PANDA’s motif filter retains reads or clones containing all requested motifs, entered directly or supplied one per line in a text file. The filter uses exact matching for predefined haplotypes; approximate matching is not implemented. Reads with errors within the motif may therefore be excluded. Motifs should be specified as they are expected to appear in bisulfite-converted reads. Non-C/T polymorphisms are preferable because C/T variation may be confounded with cytosine conversion and methylation status. Motifs should be long enough to distinguish the intended allele, while unnecessary flanking bases increase the chance that an error prevents an exact match.

### Bisulfite-aware alignment, methylation calling, and strand detection

PANDA performs reference-coordinate methylation calling without a genome-wide mapper. The reference amplicon and each input sequence are converted in silico by replacing cytosine (C) with thymine (T), then aligned through the pwalign R package using type = “global-local”, with match = 1, mismatch = –3, gap opening = –10, and gap extension = –4. Parallelization is restricted to independent read-level computations and does not alter alignment scoring, strand selection, filtering, coordinate mapping, or methylation calls.

The selected alignment is mapped back to the original, unconverted reference. CpG positions are defined by scanning the unconverted reference for CG dinucleotides before alignment. At each reference CpG, an aligned read of C is interpreted as methylated and an aligned read of T as unmethylated, subject to the read-level filtering thresholds. Exported tables and visualizations retain the original reference coordinates rather than the display order of a plot.

Strand orientation is evaluated using up to ten representative sequences and non-CpG conversion rates. The reverse-complement reference is selected only when its conversion rate exceeds 50% and is more than 20 percentage points higher than that of the forward orientation. The ten-sequence cap limits diagnostic alignments. The conversion cutoff and orientation margin are heuristic settings that require a detectable conversion signal and a difference between orientations; their sensitivity was evaluated in the controlled benchmarks. When orientation is uncertain, PANDA reports a low-confidence warning and diagnostic scores, and the reference orientation can be set explicitly. This changes the coordinate-mapping orientation without inferring unobserved sequence or altering methylation calls.

### Dereplication, quality control, paired-end handling, and display limits

To avoid aligning identical reads repeatedly, PANDA collapses NGS reads that are identical across the full retained sequence into one sequence variant. Count records the number of retained reads represented by each variant. Alignment and CpG calling are performed once per variant, whereas downstream calculations use Count, giving a variant observed 100 times the same numerical contribution as 100 identical reads analyzed separately. Sanger sequences are analyzed individually with Count = 1. By default, all retained reads are analyzed; max_reads, max_unique_reads, and min_count act as explicit computational or multiplicity filters and should be reported when used.

Phred scores are not used as PANDA’s primary retention criterion, permitting a common downstream representation for Sanger and NGS inputs. FASTQ quality assessment and adapter/primer trimming should nevertheless be performed with tools such as Trimmomatic (Bolger et al., 2014) or fastp (Chen et al., 2018; Chen, 2023), with settings recorded. Default thresholds are identity ≥ 90% and non-CpG conversion ≥ 95%. Overlapping paired-end reads should generally be merged before analysis; unmerged mode links R1 and R2 by pair identity after mapping both ends to the same reference and is intended for long amplicons with an unsequenced internal interval.

For NGS lollipop plots, users can limit the display to a specified number of the most abundant retained sequence variants (30 by default). This display limit is referred to as “Top-N” in the interface. Variants are selected by retained read count and then arranged by mean methylation level to facilitate comparison of methylation states. The adjacent bars show the retained read count corresponding to each displayed variant. This selection and ordering are used only for visualization and do not affect analytical tables, methylation estimates, heterogeneity metrics, group comparisons, or complete sequence-level exports. Binned abundance heatmaps and methylation distributions are generated from all retained variants using their Count values; heatmap binning changes only the graphical resolution.

### Statistical framework and heterogeneity metrics

PANDA reports three amplicon-adapted heterogeneity metrics: Amplicon PDR (proportion of discordant reads), Window Epipolymorphism, and Amplicon qFDRP (quantitative fraction of discordant read pairs), informed by the benchmarked framework (Scherer et al., 2020) and the original PDR formulation (Landau et al., 2014). In the definitions below, w(i) denotes the Count assigned above: w(i) = 1 for a Sanger clone and w(i) = the retained read count for a dereplicated NGS sequence variant. Count can include PCR amplification and is therefore interpreted as retained read abundance rather than the number of original biological molecules. Each metric is returned with an eligibility status and the relevant numbers of records, windows, or read pairs, distinguishing a value of zero from a non-estimable result.

For Amplicon PDR, records covering fewer than four CpGs were excluded. Each eligible record i was classified as discordant [D(i) = 1] when it contained both methylated and unmethylated CpG calls, and as concordant [D(i) = 0] otherwise. Amplicon PDR is the abundance-weighted proportion of eligible records classified as discordant: the weights of discordant records are summed and divided by the total weight of all eligible records. Amplicon PDR was calculated as:

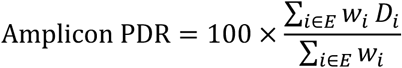

For each window v of four consecutive CpGs, records with complete calls were assigned to one of the 16 possible binary methylation patterns. Each record contributed its weight w(i) to the count of its observed pattern, and the abundance-weighted pattern frequency p(v,k) was obtained by dividing that weighted count by the total eligible weight in the window. Windows with weighted coverage below two were excluded by default. Window-level Epipolymorphism and the amplicon-level arithmetic mean across eligible windows were calculated as:

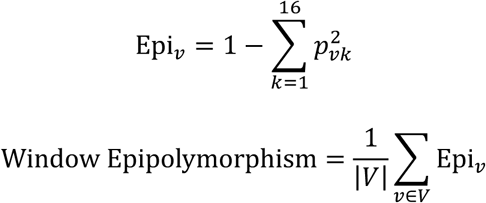

The window-level value is the probability that two independently sampled epialleles from the same window have different four-CpG patterns. PANDA reports the unweighted arithmetic mean across eligible windows and exports each window’s coverage and score.

Amplicon qFDRP describes the average methylation disagreement expected between two retained reads or Sanger clones. For each pair of sequence records i and j, S(i,j) comprised the CpGs observed in both records, and m(i,c) represented the binary methylation call for record i at CpG c. The proportion of discordant calls among the shared CpGs was calculated as the normalized Hamming distance d(i,j):

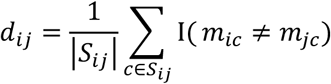

Pairs sharing fewer than four CpGs were excluded by default and the remaining pairs formed the eligible set P. Distances were calculated over all CpGs shared by each eligible pair; this minimum requirement is an eligibility threshold, not a window width. Two distinct NGS variants i and j represent w(i)w(j) possible read pairs and therefore contribute that pair weight. A variant with weight w(i) also represents w(i)[w(i) – 1]/2 pairs of reads drawn from the same variant; these pairs are retained with distance zero because their methylation patterns are identical. Sanger clones have weight one and consequently contribute only pairs between distinct clones. Amplicon qFDRP was calculated as:

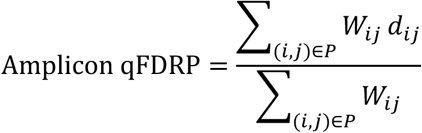

This weighting gives the same qFDRP as first expanding every dereplicated NGS variant into Count identical reads and then evaluating all eligible read pairs, but avoids constructing the expanded dataset. Same-variant pairs contribute zero to the numerator and remain in the denominator because they are part of the retained read population. PANDA exports weighted eligible-pair and eligible variant-pair counts, the weighted median shared-CpG count, and the shared-CpG distribution. None of the three metrics is interpreted as a standalone test for ASM.

### Read-level outputs and group comparison

After filtering and alignment, PANDA records CpG calls for each retained Sanger clone or dereplicated NGS sequence variant in reference coordinates. These calls support phased lollipop plots, read-count-aware binned heatmaps, methylation distributions, ASM-oriented clustering, heterogeneity summaries, and alignment and quality-control tables. The GUI presents these results interactively, whereas the CLI exports the corresponding tables, plotting data, and PDF figures for scripted analysis.

For multi-sample analyses, PANDA first summarizes each sample and then supports predefined two-group comparisons. Overall methylation is compared between independent samples using a two-sided Wilcoxon rank-sum test when each group contains at least two samples; individual reads are not pooled for this test. With its default settings, stats::wilcox.test() calculates an exact *p*-value when the sample sizes and absence of ties permit; otherwise, it uses a normal approximation with continuity correction.

At each CpG, PANDA applies a two-sided Welch’s *t*-test when at least two finite sample-level methylation estimates are available in each group. A comparison is reported as not estimable when coverage is insufficient or the test cannot be calculated, for example when both groups have zero variance. In such cases, nominal and adjusted *p*-values are reported as NA. No pooled read-level test is substituted because independent samples, rather than individual reads, are the units of inference. The output records the numbers of finite sample-level estimates in each group (N_Samples_1 and N_Samples_2) and the reason for non-estimability (Test_Status). Benjamini–Hochberg adjustment is applied only to estimable CpGs within each comparison.

### Synthetic demonstration datasets and analyses

Sanger and amplicon-NGS datasets were generated with the PANDA_demo_data_generation.R script, available in the project repository. Mouse target sequences were extracted from the *Mus musculus* mm10 reference genome using Biostrings (version 2.80.1) and BSgenome.Mmusculus.UCSC.mm10 (version 1.4.3) R packages. Three replicates were generated per condition, with 3,000 reads per NGS sample and 16 clones per Sanger sample.

Experiment A used the *H19* amplicon to compare an ASM-like state containing equal proportions of low- and high-methylation populations (assigned mean probabilities of 0.02 and 0.98) with an LOI-like state assigned a probability of 0.98 throughout. Experiment B comprised a 1:1 mixture of two *H19* haplotypes distinguished by an engineered central A/G variant and assigned methylation probabilities of 0.02 and 0.98, respectively. Haplotype selection was evaluated using 21-bp motifs spanning the variant. Experiment C used the *Nanog* amplicon with contrasting wild-type and knockout profiles. Low methylation was assigned to CpGs at amplicon positions 80, 106, 116, 177, 214, and 248 in wild-type and positions 1, 5, and 77 in knockout; the remaining CpGs were assigned high methylation.

For each read and CpG, a methylation probability was independently sampled from a beta distribution with shape parameters equal to 50 times the assigned mean and 50 times its complement. A methylated or unmethylated call was then sampled from a Bernoulli distribution using that probability. Non-CpG cytosines were converted to thymine, and per-base substitution errors were introduced at rates of 0.005 for NGS and 0.001 for Sanger sequencing. The random seed was set to 11.

The demonstrations evaluated global methylation estimates, read-level methylation patterns, motif-based haplotype selection, and within-sample heterogeneity. For each sample, the expected global methylation level was defined as the mean of the sampled CpG probabilities before methylated or unmethylated calls were generated. Agreement with the PANDA estimate was summarized separately for NGS and Sanger data using signed error, mean absolute error (MAE), and root mean square error (RMSE). Signed error was calculated as the PANDA estimate minus the expected value. For motif-filter evaluation, retained reads carrying the intended haplotype were classified as true positives, retained reads carrying the other haplotype as false positives, and excluded reads carrying the intended haplotype as false negatives. Sensitivity and positive predictive value were calculated from these counts.

### Controlled computational benchmarks

70 synthetic benchmark datasets were generated with seed 11 from GRCm38/mm10 amplicons at *H19*, *Meg3*, and *Nanog*. Source-molecule identities, CpG states, and reference coordinates were recorded as truth. Unless otherwise specified, PANDAcore v1.1.0 used minimum alignment identity of 90% and minimum non-CpG conversion of 95%. The datasets, the R scripts used to generate them, and documentation describing their contents are provided in Data S1.

To quantify how incomplete conversion and substitution errors affect read retention, CpG calls, and haplotype assignment, fixed H19-based molecule draws were perturbed in three replicates of 1,200 reads sampled from 300 molecules. Conversion probabilities of 100%, 99%, 97.5%, 95%, and 90% were crossed with substitution rates of 0%, 0.5%, and 2%. Read retention, retained-read CpG-call accuracy, and methylation error against pre-QC read truth were measured. Engineered A/G haplotypes were used to score motif assignment, comparing exact matching with a one-flanking-mismatch rule at 9, 13, 17, or 21 bp; the central allele had to match under both rules.

To test whether PDR, Window Epipolymorphism, and qFDRP distinguish patterns that have similar site-wise means, defined methylation architectures were generated. Four Meg3-based populations comprised 400 molecules and 1,200 sampled reads each, with approximately 50% mean methylation but bimodal, block, checkerboard, or independent CpG states. A spatially patterned Nanog population provided an additional configuration. Metric values were compared for original molecules, sampled reads, and PANDA output.

Unequal molecule recovery was examined to distinguish changes in read counts from changes in the source population. From 320 *H19*-based molecules carrying 16 complementary patterns, 600 reads were selected with equal or Gamma-distributed (shape = 2, rate = 2) weights, or with weights producing 1.5:1 and 1:1.5 selection ratios for fully methylated versus unmethylated molecules. Each condition was generated with 0% or 0.5% post-sampling substitution error. This modeled differential read yield without simulating PCR cycles. PANDA’s count-weighted and equal-unique-variant-weighted estimates were compared with molecule and sampled-read truth.

Strand assignment was evaluated in five *Meg3*-based unmerged-pair datasets with reversed references, 20% or 50% contamination, substitution errors, or 75% conversion. Sensitivity was assessed by varying the number of representative sequences (5–50), conversion cutoff (40–60%), and orientation margin (10–30 percentage points), using the non-CpG-only scoring in PANDAcore v1.1.0.

Scaling was assessed in seven *H19*-based datasets. At 20,000 reads, the number of exact distinct input sequence patterns was set to 500, 1,000, 3,000, 5,000, or 10,000; two additional datasets held this number at 3,000 with 10,000 or 50,000 reads. Single-worker PANDA CLI wall time, including R startup, and peak resident memory were measured using R 4.6.1 on macOS 15.7.3 with an Intel Xeon W-3223 CPU (3.50 GHz) and 192 GiB RAM.

### Re-analysis of public *ST8SIA1* targeted bisulfite sequencing data

Public paired-end TBAS data from the *ST8SIA1* locus in three chimpanzee and three rhesus macaque placentae samples (Bogutz et al., 2019) were re-analyzed using PANDA. The chimpanzee reference amplicon was 320 bp, whereas the paired FASTQ reads were 151 bp; the combined nominal read length was therefore 18 bp shorter than the amplicon. R1 reads aligned in both orientations relative to the amplicon reference. Chimpanzee R1 reads were first analyzed alone as an orientation-dependent single- end control and then together with their matched R2 reads in unmerged mode; overlap-based merging was not applied to these samples. The rhesus macaque samples were analyzed using fastp-merged FASTQ files. No max_reads or max_unique_reads limits were applied. Minimum alignment identity, non-CpG conversion, read count per unique NGS sequence, shared CpGs for Amplicon qFDRP, and four-CpG-window coverage were set to 90%, 95%, 1, 4, and 2, respectively. In the chimpanzee unmerged analysis, matched R1 and R2 reads were independently aligned to the same reference coordinate system and combined by pair identity, with positions in the intervening unsequenced interval retained as missing.

### Software availability and verification

PANDA is distributed under the MIT License. All analyses reported in this study used PANDA and PANDAcore v1.1.0. The source code, installation instructions, documentation, synthetic demonstration data, and code for PANDAcore tests are available at https://github.com/kubo-azu/PANDA. An optional hosted Shiny GUI is available at https://huggingface.co/spaces/kubo-azu/PANDA. Local installation is supported on macOS and Linux, with WSL recommended for Windows.

The package versions recorded in `renv.lock`, the example analysis configurations, and machine-readable parameter metadata support reproducibility. Automated tests evaluate core PANDAcore functions, heterogeneity metrics, and the equivalence of serial and parallel calculations.

## Results

### Demonstration using synthetic bisulfite amplicon datasets

Synthetic NGS and Sanger datasets with predefined methylation and sequence-defined haplotype states were analyzed using the PANDA workflow. Across the synthetic demonstrations, all input reads and clones were retained under the specified alignment identity and non-CpG conversion thresholds. Overall methylation and all heterogeneity metrics were estimable in every sample-level analysis (Data S2).

#### 1. Visualization of epiallele architectures and methylation patterns

The synthetic datasets were processed through the same computational workflow used by the GUI and CLI. Ranked lollipop plots displayed the 30 most abundant variants, selected by retained read count and ordered by mean methylation state, while binned abundance heatmaps summarized all retained reads while preserving physical CpG coordinates. The ASM-like *H19* sample showed two abundant groups of predominantly methylated and predominantly unmethylated patterns (Figure 2A, C). In contrast, the LOI-like sample was dominated by highly methylated patterns (Figure 2B, D). The corresponding all-retained-read methylation distributions are shown in Figure 2E and F.

**Figure 2.**
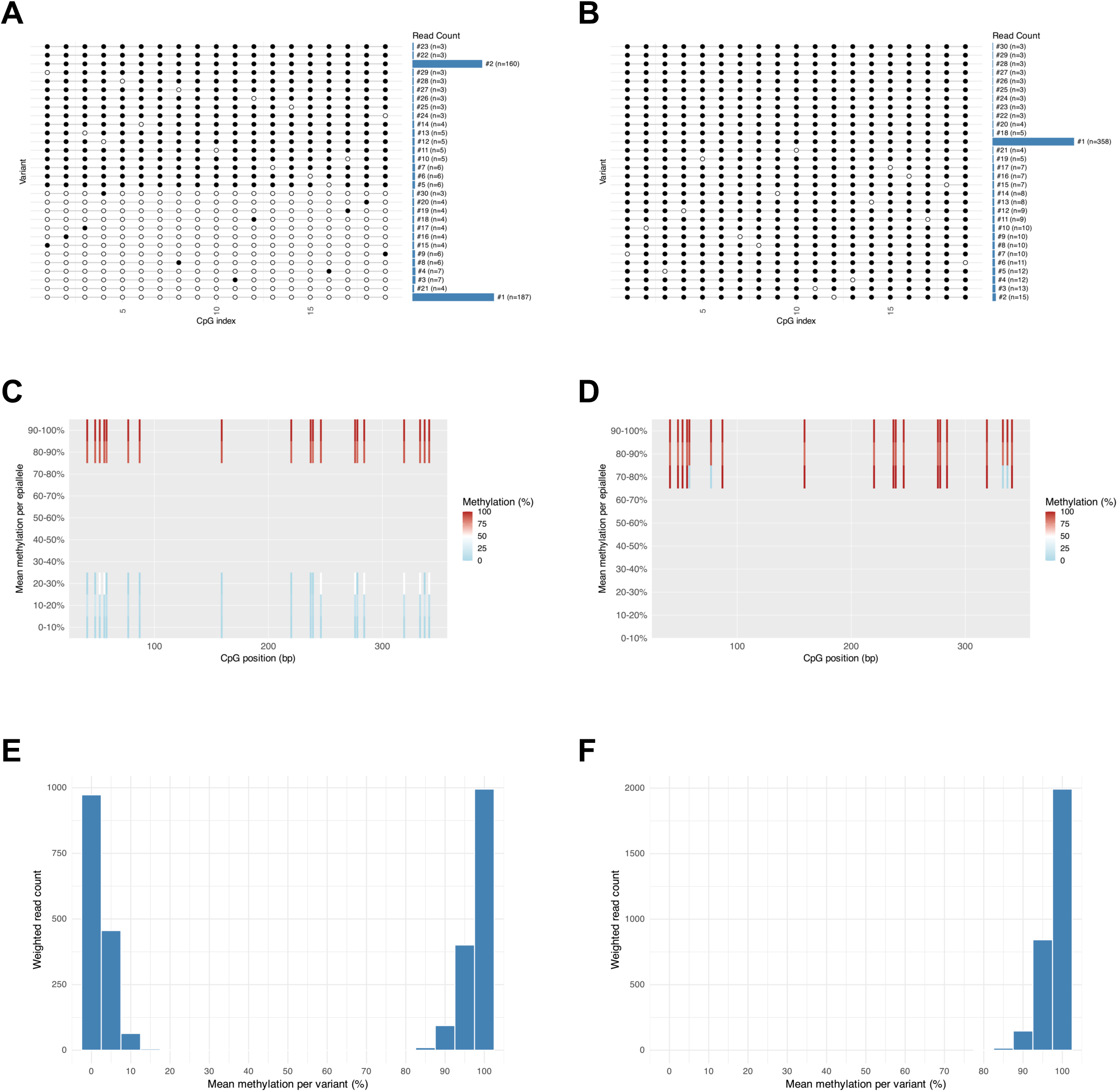
Read-level visualization of structured ASM-like and LOI-like methylation states in synthetic amplicon-NGS data. **(A,B)** Ranked lollipop plots for representative Experiment A ASM-like **(A)** and LOI-like *H19* **(B)** samples, respectively. The plots show the 30 most abundant sequence variants retained after filtering, with horizontal bars representing their read counts. Filled and open circles indicate methylated and unmethylated CpGs, respectively. **(C,D)** Binned abundance heatmaps for the same ASM-like **(C)** and LOI-like **(D)** samples, respectively, constructed from all retained reads and shown against physical CpG coordinates. Vertical extent represents retained read abundance, and color represents methylation state. **(E,F)** Read-count-weighted methylation distributions for the same ASM-like **(E)** and LOI-like **(F)** samples, respectively, calculated using all retained reads.

#### 2. Global methylation estimates in the demonstration datasets

PANDA estimates were close to the simulation-derived expected methylation values (Figure S1, Data S3). For unfiltered NGS data, MAE was 0.092, 0.039, and 0.087 percentage points in Experiments A–C, respectively, with corresponding RMSE values of 0.120, 0.042, and 0.096 percentage points. For Sanger data, MAE was 0.409, 0.731, and 0.788 percentage points in Experiments A–C, respectively.

#### 3. Motif-based haplotype filtering

In the mixed Experiment B NGS datasets, exact-match motifs selected reads from the two sequence-defined haplotypes with sensitivities of 90.07% and 90.36% and positive predictive values of 99.85% and 99.80%, respectively (Data S3). Mean methylation was 50.00% before filtering and 2.31% or 97.65% after selection for motif A or B, respectively. Thus, motif filtering recovered the methylation states assigned to the two haplotypes. The reduced sensitivity was consistent with simulated substitution errors disrupting exact matches within the motifs.

#### 4. Quantitative assessment of within-sample heterogeneity

The three heterogeneity metrics responded differently to the predefined methylation structures. In Experiment A, PDR was nearly identical between the NGS ASM and LOI samples (33.90% and 34.02%, respectively; Figure 3A). By contrast, Window Epipolymorphism declined from 0.579 to 0.158 (Figure 3B), and Amplicon qFDRP from 0.500 to 0.042 (Figure 3C). Experiment B showed a comparable response to haplotype selection. The unfiltered mixture yielded a Window Epipolymorphism of 0.579 and qFDRP of 0.500, whereas selection for either motif reduced these values to approximately 0.162 and 0.045, respectively. In Experiment C, the wild-type and knockout groups produced similar values for all three metrics. PDR was 100% in both groups, while Window Epipolymorphism was 0.159 and 0.163 and qFDRP was 0.042 and 0.044, respectively.

**Figure 3.**
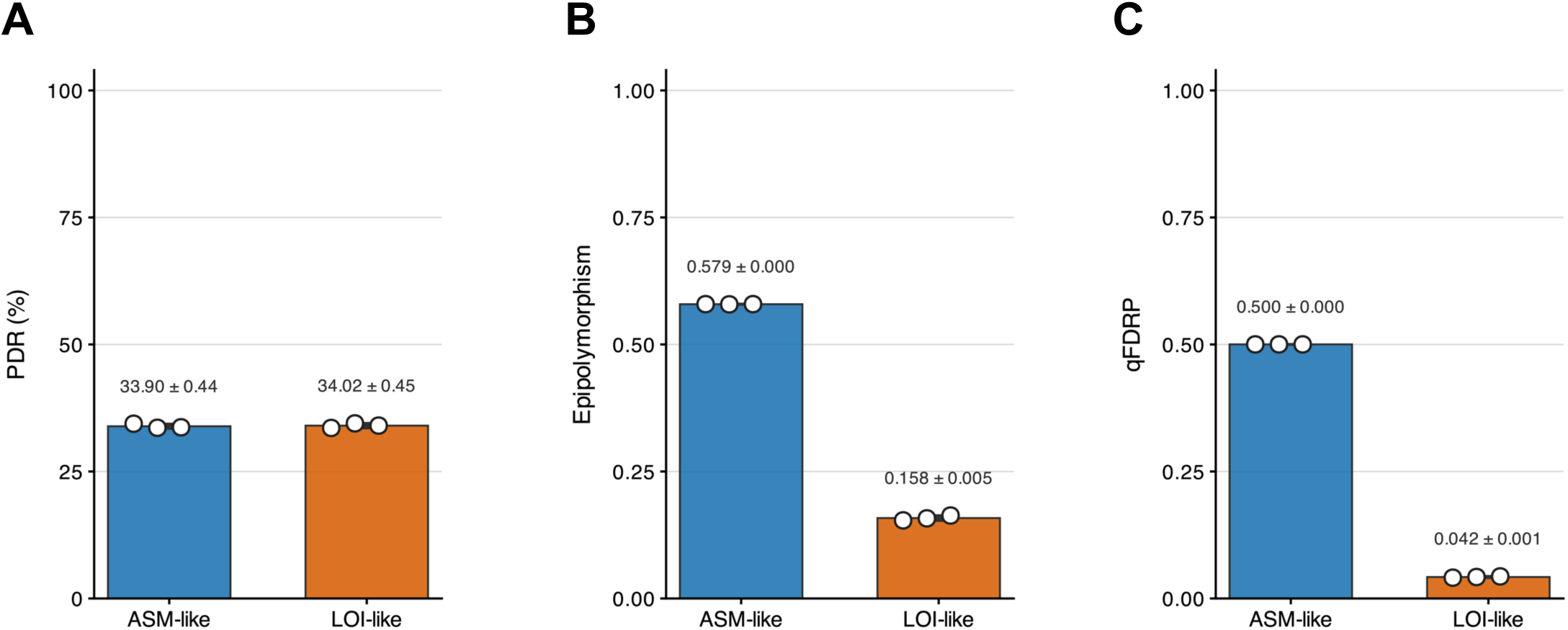
PANDA heterogeneity metrics in synthetic Experiment A amplicon-NGS datasets. Replicate-level values are shown for Amplicon PDR **(A)**, Window Epipolymorphism **(B)**, and Amplicon qFDRP **(C)** in the ASM-like and LOI-like states. Bars represent group means, error bars represent standard deviations, and white points represent independently generated synthetic replicates, not biological replicates.

Corresponding Sanger analyses were based on 16 clones per replicate, with Count = 1 for each clone. The heterogeneity estimates showed the same qualitative relationships among the predefined methylation states (Figure S2). Representative Sanger methylation patterns are shown in Figure S3.

#### 5. Recovery of group-level methylation differences

Group comparisons reproduced the intended global and CpG-specific differences between the synthetic conditions (Figure 4A, B). In Experiment A, mean NGS methylation was 49.97% in the ASM group and 97.84% in the LOI group. In Experiment C, the corresponding values were 45.70% in the wild-type group and 71.72% in the knockout group. Each group comprised three independently generated samples. The exact Wilcoxon rank-sum test returned *p* = 0.10 for both global comparisons. Given the small number and synthetic nature of the replicates, these tests demonstrate the group-comparison workflow rather than provide biological inference.

**Figure 4.**
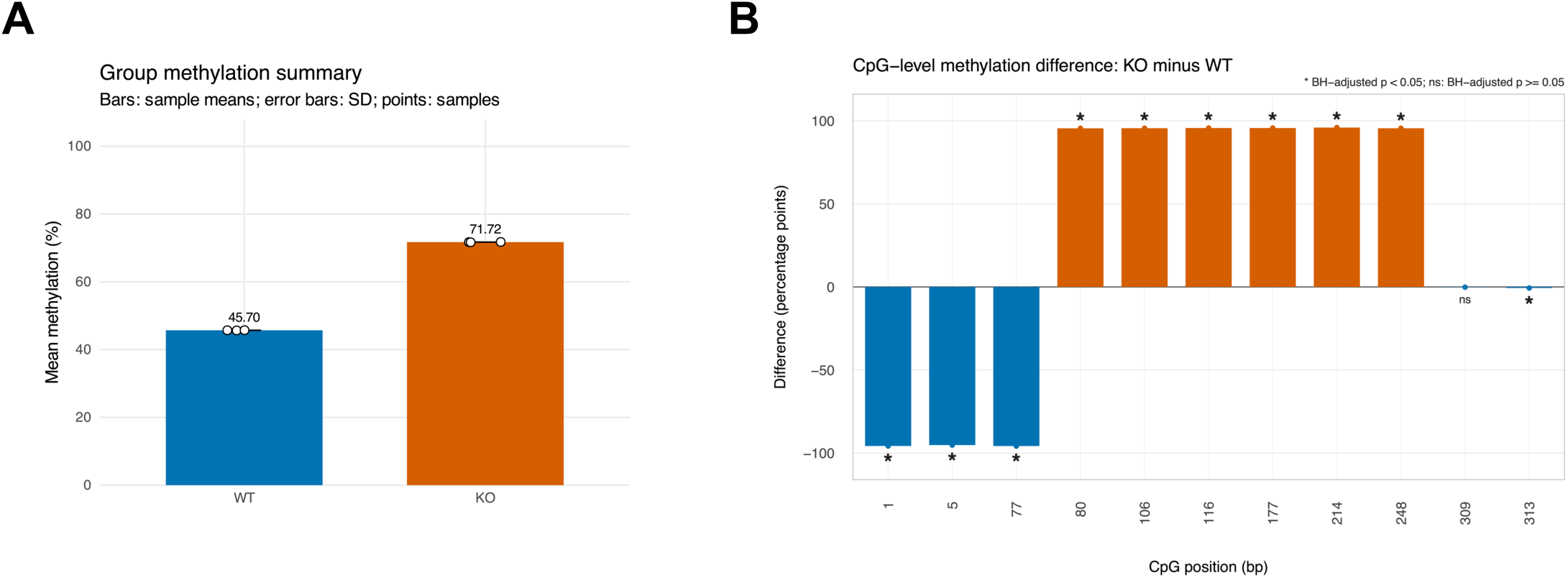
Overall and CpG-level group comparison in synthetic Experiment C amplicon-NGS data. **(A)** Overall methylation in three independently generated synthetic replicates per group. Bars show group means, error bars show standard deviations, and points show individual replicates. **(B)** Difference in pooled CpG methylation percentages (knockout minus wild-type, in percentage points), with CpGs arranged in reference-position order. Asterisks indicate Benjamini–Hochberg-adjusted *p* < 0.05 based on replicate-level comparisons; “ns” indicates adjusted *p* ≥ 0.05.

At the CpG level, all 19 NGS positions in Experiment A and all 11 positions in Experiment C were estimable. After Benjamini–Hochberg adjustment, all 19 Experiment A positions and all nine Experiment C positions assigned different methylation probabilities had adjusted *p*-values below 0.05; one of the two Experiment C positions assigned the same probability in both groups (position 313) also met this threshold. In the Sanger data, 15 of 19 Experiment A positions and 7 of 11 Experiment C positions met this threshold. Four Experiment A positions and three Experiment C positions had zero variance in both Sanger groups and were therefore reported as not estimable. Complete CpG-level results, including test status, coverage, and effect direction, are provided in Data S3.

### Evaluation using controlled synthetic benchmarks

As conversion probability decreased, QC retention fell more sharply than CpG-call accuracy among reads that passed QC. At a 0.5% substitution rate, lowering conversion from 100% to 95% reduced mean on-target QC recall from 100% to 51.7% (Figure 5A), while retained-read CpG-call accuracy declined from 99.8% to 97.6% (Figure 5B). Relative to the full pre-QC read population, CpG-wise methylation MAE increased from 0.10 to 3.15 percentage points, and correct exact-motif haplotype assignment declined from 90.4% to 67.2% (Table S2). At 90% conversion, only 7.0% of reads passed QC; the 94.7% CpG-call accuracy was estimated from this retained subset (Figure 5A,B).

**Figure 5.**
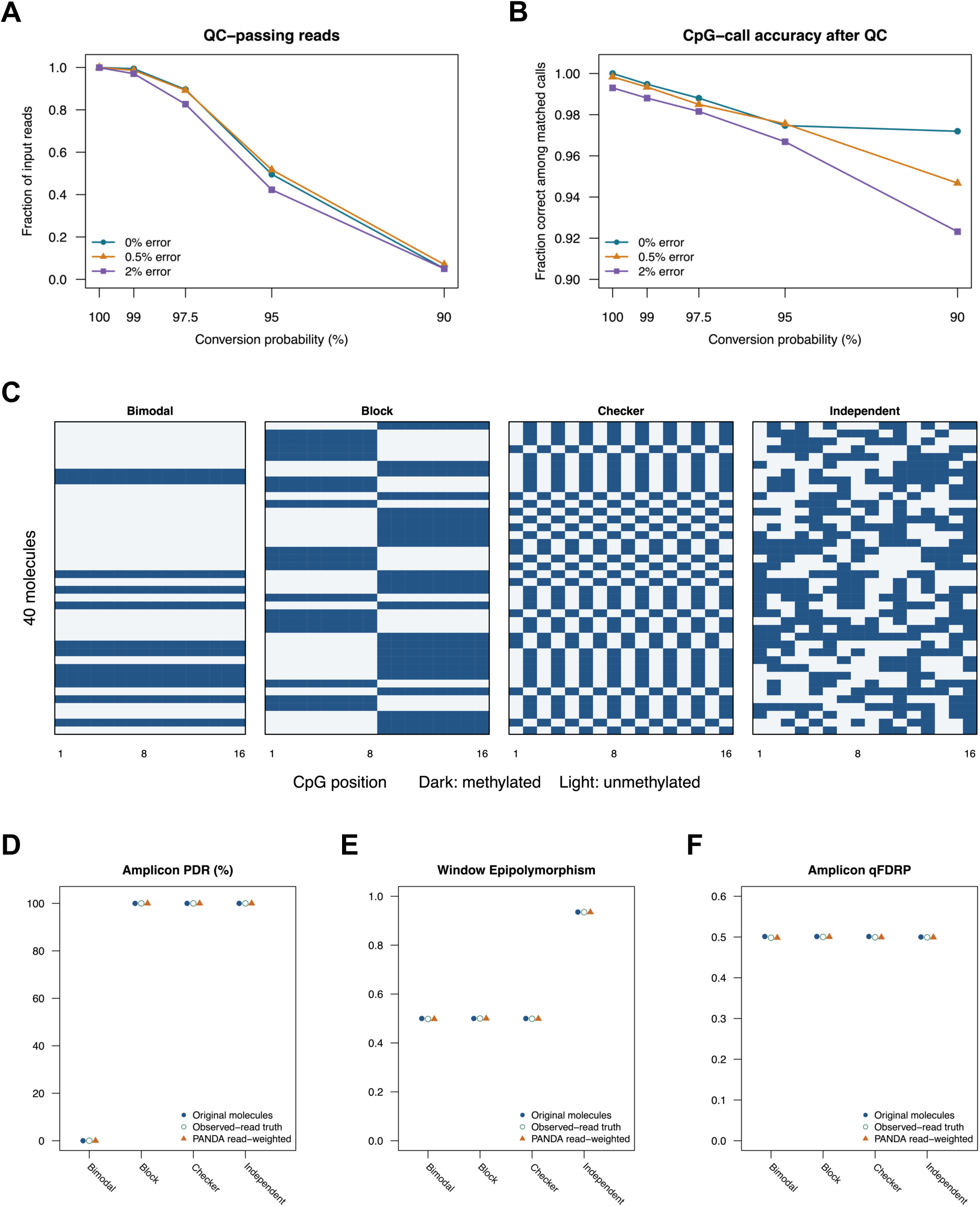
Read retention, CpG-call accuracy, and heterogeneity in controlled synthetic benchmarks. **(A,B)** On-target QC recall (**A**; fraction of input reads passing QC) and CpG-call accuracy among retained matched calls **(B)** across five conversion probabilities and three per-base substitution rates in *H19*-based datasets. Points show means of three independently generated synthetic replicates per condition; lines distinguish substitution rates. **(C)** Methylation states at the same 16 CpGs for the first 40 original molecules in each of four *Meg3*-based populations: bimodal, block, checkerboard, and independent. Dark cells indicate methylated CpGs and light cells indicate unmethylated CpGs. **(D–F)** Amplicon PDR **(D)**, Window Epipolymorphism **(E)**, and Amplicon qFDRP **(F)** for these four populations. Filled blue circles show metrics calculated from 400 original molecules, open green circles show truth for 1,200 sampled reads, and orange triangles show PANDA’s read-weighted estimates.

At 100% conversion and 0.5% substitution error, correct motif assignment by exact matching declined from 95.7% with a 9-bp motif to 90.4% with a 21-bp motif. Allowing one flanking mismatch increased correct assignment to 99.1% and 99.0%, respectively, while allele specificity remained above 99% in this simulation (Table S3).

Populations with approximately 50% mean methylation showed distinct arrangements of methylated CpGs across molecules (Figure 5C). PDR separated the bimodal pattern (0%) from the block, checkerboard, and independent patterns (100%; Figure 5D). Window Epipolymorphism was approximately 0.50 for the first three patterns and 0.94 for the independent pattern (Figure 5E), whereas qFDRP remained approximately 0.50 for all four (Figure 5F). In these datasets, all reads covered the same CpG set without missing calls; qFDRP was therefore determined by the CpG-wise methylation frequencies and retained read count and did not distinguish the different linkage patterns. For each metric, PANDA’s read-weighted estimate matched the truth calculated from sampled reads (Figure 5D–F).

Changing molecule-selection weights altered the sampled-read population while leaving the original molecule population fixed (Figure S4). In the error-free conditions, count-weighted estimates matched sampled-read truth under all four weighting schemes. Preferential recovery of fully methylated or unmethylated molecules yielded mean methylation of 52.5% or 46.6% among sampled reads, compared with 50.0% among original molecules. Sampled-read PDR was 89.2% in both conditions, compared with 87.5% in the original population. Equal weighting of distinct variants yielded PDR of 87.5% but qFDRP of 0.533, compared with original-molecule qFDRP of 0.502. With 0.5% substitution error, count-weighted PDR differed from sampled-read truth by at most 0.5 percentage points.

Automatic reference-orientation detection was correct in all five contamination, reference-reversal, and low-conversion scenarios when the default ten representative read ends were used. This held across the tested conversion cutoffs (40–60%) and orientation margins (10–30 percentage points). With 50 read ends, the reversed-reference scenario containing 50% contaminants was assigned incorrectly and flagged as low confidence. The 75%-conversion scenario was oriented correctly but yielded no reads passing the separate 95% non-CpG conversion QC filter.

At 20,000 reads, wall time increased from 14 s for 500 distinct input sequences to 24 s for 1,000, 101 s for 3,000, 258 s for 5,000, and 1,070 s for 10,000 (Figure S5). Peak resident memory across these five conditions was 635–1,270 MB. At fixed 3,000 distinct sequences, 10,000, 20,000, and 50,000 reads required 104, 101, and 106 s, respectively; peak resident memory ranged from 701 to 792 MB.

### Re-analysis of public TBAS datasets from primate placentae

The *ST8SIA1* locus re-analysis included three chimpanzee and three rhesus macaque placental datasets (Bogutz et al., 2019). Chimpanzee libraries contained 1,091–1,761 input read pairs per sample and were analyzed both as R1-only controls and as coordinate-linked unmerged R1/R2 pairs. The fastp-merged rhesus macaque libraries contained 1,073–1,431 reads per sample.

In all three chimpanzee datasets, retained R1 reads aligned in both orientations relative to the amplicon reference. Consequently, an individual R1-only profile represented one of two possible read ends rather than a fixed side of the amplicon (Figure 6A). R1-only and coordinate-linked R1/R2 profiles are compared in Figure 6B and C. Mean overall methylation was 46.37% (SD 3.96%) in the R1-only analyses and 46.10% (SD 4.23%) after coordinate linkage. The corresponding Amplicon qFDRP values were 0.495 (SD 0.005) and 0.494 (SD 0.008), respectively. In contrast, mean Amplicon PDR increased from 37.08% to 56.23%, and the median number of CpGs shared by eligible qFDRP pairs increased from 9 to 21 after coordinate linkage (Table S4).

**Figure 6.**
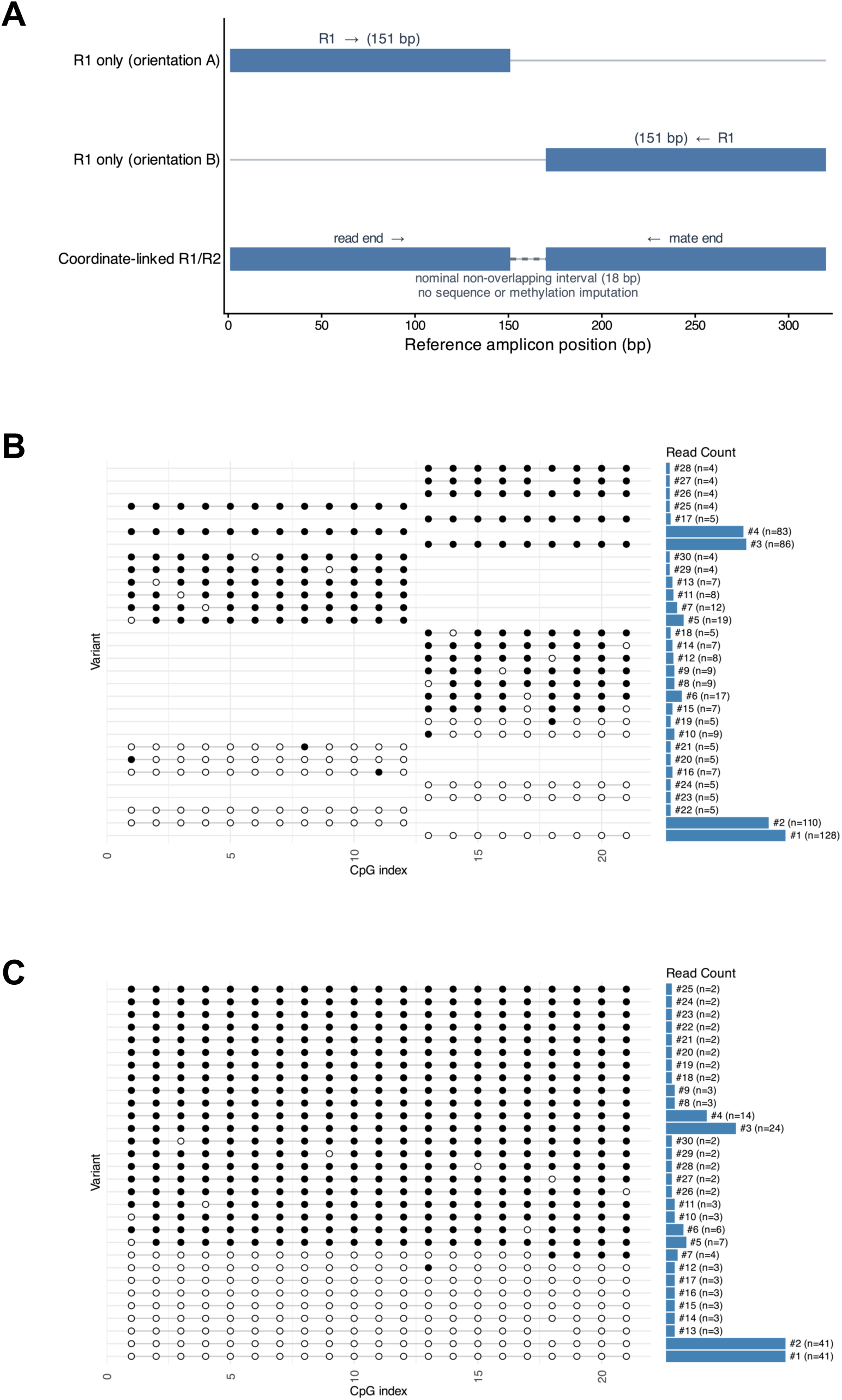
Coordinate-linked re-analysis of paired-end *ST8SIA1* reads from chimpanzee placentae. **(A)** Schematic of the nominal observation geometry for the 320-bp chimpanzee reference amplicon and 151-bp paired-end reads. **(B,C)** Ranked lollipop plots comparing R1-only analysis **(B)** with coordinate-linked R1/R2 analysis **(C)** in a representative chimpanzee sample. Filled and open circles denote methylated and unmethylated CpGs, respectively, and horizontal bars show retained read counts for the 30 most abundant sequence variants. Note that variants are ranked independently in the two plots; therefore, rows at the same vertical position do not necessarily represent the same sequence.

The coordinate-linked chimpanzee samples showed abundant read populations at both low and high methylation levels (Figure 7A, B). Across the three samples, mean overall methylation was 46.10% (SD 4.23%), Amplicon PDR was 56.23% (SD 5.41%), Window Epipolymorphism was 0.593 (SD 0.016), and Amplicon qFDRP was 0.494 (SD 0.008). In contrast, the three merged rhesus macaque samples were dominated by reads with little or no methylation (Figure 7C, D), with mean overall methylation of 0.72% (SD 0.13%), Amplicon PDR of 8.90% (SD 1.58%), Window Epipolymorphism of 0.0586 (SD 0.0079), and Amplicon qFDRP of 0.0144 (SD 0.0026). The abundance heatmaps of the remaining four samples are provided in Figure S6.

**Figure 7.**
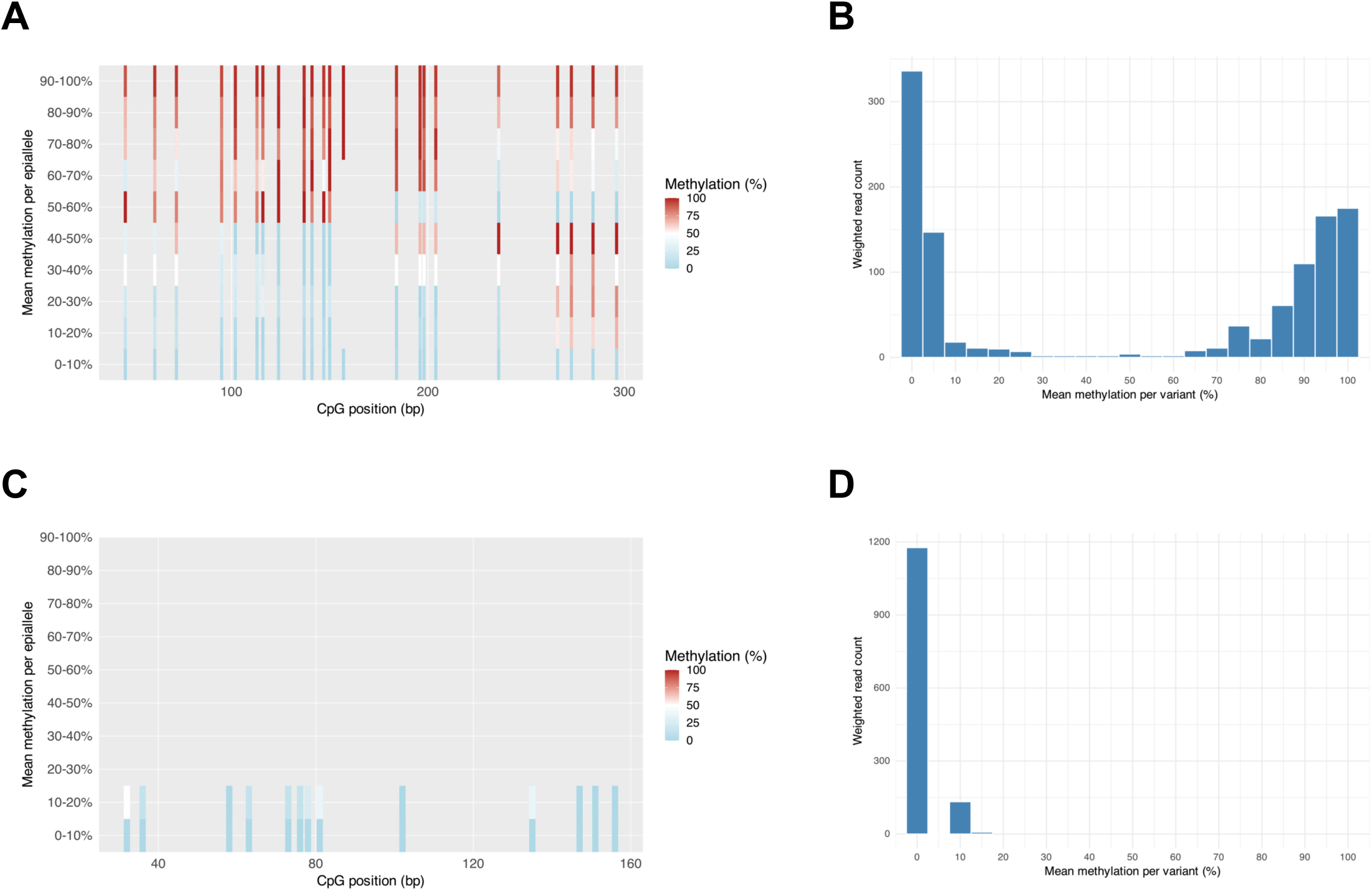
Species-associated *ST8SIA1* locus methylation patterns in chimpanzee and rhesus macaque placentae datasets. **(A)** Binned abundance heatmap for the representative coordinate-linked chimpanzee sample 20-c1_S43_L001, constructed from all retained read pairs and plotted at physical CpG coordinates. **(B)** Read-count-weighted distribution of mean methylation per retained sequence variant for the same sample. **(C)** Binned abundance heatmap for the representative merged rhesus macaque sample 42-r1_S92_L001, constructed from all retained reads and plotted at physical CpG coordinates. **(D)** Read-count-weighted methylation distribution for the same macaque sample.

## Discussion

PANDA provides a workflow for phased, read-level analysis of targeted bisulfite amplicon NGS, with additional support for Sanger-derived sequences. It preserves CpG methylation patterns in physical reference coordinates and incorporates observed NGS read abundance, allowing visualization, heterogeneity assessment, haplotype filtering, and group comparison within a common analytical representation. The synthetic datasets illustrated methylation profiling, sequence-defined haplotype selection, group comparison, and heterogeneity summaries, while the public *ST8SIA1* locus re-analysis demonstrated coordinate-linked analysis of unmerged paired reads.

The synthetic demonstration datasets were designed to illustrate software functions under controlled conditions. Global methylation estimates were close to the simulation-derived expected values, and exact-match motif selection recovered the intended haplotype-associated methylation contrast. Sensitivities of approximately 90% and positive predictive values above 99.8% describe the specified motif lengths and substitution-error model; reads with errors within the motif could fail selection. The motif-length sensitivity analysis showed that longer exact-match motifs lost more reads, whereas allowing one flanking mismatch improved recovery in the simulated construct. Because PANDA uses exact matching, motif length should be limited to what is needed to identify the intended allele.

The controlled conversion and error benchmarks distinguished accuracy among retained reads from representation of the input population. At a simulated conversion probability of 95% and 0.5% substitution error, only 51.7% of on-target reads passed the default QC criteria, although CpG-call accuracy among retained reads was 97.6% (Figure 5A,B). Thus, high retained-read accuracy can coexist with substantial read loss and a shift in methylation estimates relative to the pre-QC population. These simulations do not determine an experimental conversion cutoff.

The three metrics describe different aspects of the observed read population and are most informative when interpreted alongside CpG-wise methylation. Amplicon PDR quantifies the prevalence of reads containing both methylated and unmethylated CpGs. In the matched-mean benchmark, it separated a bimodal mixture of uniformly methylated and unmethylated molecules (PDR 0%) from populations composed of internally mixed reads (100%; Figure 5D). Thus, PDR helps distinguish variation between reads from mixed methylation within reads even when CpG-wise frequencies are similar. Its binary classification explains why it was nearly identical in the ASM-like and LOI-like states (Figure 3A) and reached 100% in both Experiment C groups. Because longer CpG spans provide more opportunities for mixed calls, comparisons also require attention to read coverage (Figure 6, Table S4).

Window Epipolymorphism describes the diversity of complete four-CpG patterns within an amplicon. In the matched-mean benchmark, it was approximately 0.94 for the independent pattern but 0.50 for the bimodal, block, and checkerboard populations (Figure 5E). It therefore detected a difference in local pattern diversity that was not visible from CpG-wise frequencies or from PDR in the three populations for which it was 100%. This makes it useful when the arrangement of neighboring CpGs is of interest, provided that the windows have sufficient reads with complete calls. The score summarizes diversity rather than identifying the constituent patterns, which remain visible in the read-level plots.

Amplicon qFDRP reports the average fraction of disagreeing CpGs between eligible read pairs, weighted by retained read count. It provides a compact description of the magnitude of between-read discordance, as illustrated by its high value in the ASM-like mixture and low value in the LOI-like state (Figure 3C). This contrast is not, however, independent evidence of different CpG linkage: when reads cover the same CpGs without missing calls, qFDRP is determined by CpG-wise frequencies and read count, explaining its similar values across the four matched-mean populations (Figure 5F). With partial coverage, the set of eligible pairs and shared CpGs must also be considered. Table 2 summarizes the information provided by each metric and the conditions that affect its interpretation. Their interpretation is strengthened by phased plots, methylation distributions, coverage diagnostics, and, when assessing ASM, sequence-defined haplotypes and experimental context.

**Table 2.** Interpretive roles and limitations of heterogeneity metrics evaluated in this study.

| **Metric** | **Property** | **Interpretive value** | **Key considerations** |
|---|---|---|---|
| Amplicon PDR | Proportion of eligible reads with both methylated and unmethylated CpGs. | Distinguishes mixtures of uniformly methylated or unmethylated reads from internally mixed reads, even with similar CpG-wise frequencies. | Saturates when most reads are mixed; depends on the number and positions of observed CpGs. |
| Window Epipolymorphism | Diversity of complete four-CpG patterns across eligible windows. | Detects local pattern diversity and CpG linkage that site-wise frequencies and saturated PDR can miss. | Requires complete four-CpG calls and adequate window coverage; the score does not identify specific patterns. |
| Amplicon qFDRP | Read-count-weighted mean fraction of disagreeing CpGs between eligible read pairs. | Summarizes average between-read disagreement; can differ at the same amplicon-wide mean if CpG-wise frequencies differ. | With complete shared-CpG coverage, fully determined by CpG-wise frequencies and read count; missing calls and pair eligibility also affect it. |

PDR, proportion of discordant reads; qFDRP, quantitative fraction of discordant read pairs.

The re-analysis of TBAS datasets at the *ST8SIA1* locus was consistent with the contrasting methylation architectures previously reported for the two species (Figure 7, Table S4). Chimpanzee read-level distributions contained abundant populations at both low and high methylation levels, accounting for the intermediate mean methylation of approximately 46% and the high Amplicon qFDRP of approximately 0.49. In contrast, the rhesus macaque samples were predominantly unmethylated and showed markedly lower Window Epipolymorphism and Amplicon qFDRP. These findings illustrate why mean methylation should be interpreted together with read-level distributions and heterogeneity metrics.

Comparison of the chimpanzee R1-only and coordinate-linked R1/R2 analyses clarified the effect of paired-read linkage (Figure 6, Table S4). Overall methylation and Amplicon qFDRP were nearly unchanged, whereas the median number of shared CpGs increased from 9 to 21 and Amplicon PDR increased after coordinate linkage. The expanded CpG span provided more opportunities for an eligible record to contain both methylated and unmethylated calls, consistent with the reported sensitivity of PDR to read length (Scherer et al., 2020). Because R1 reads aligned in both orientations, R1-only analysis could retain valid reads from either end but could not recover phase between the two ends. PANDA linked matched R1 and R2 calls in reference coordinates without imputing the intervening unsequenced region. Differences in PDR between processing modes should therefore be interpreted in relation to the number and positions of CpGs observed within each record.

### Scalability and intended scope

PANDA is designed for locus-focused studies in which phased methylation patterns and within-sample heterogeneity are of interest. Its shared representation allows Sanger-derived clone sequences and amplicon-NGS reads to be analyzed using the same methylation-call and metric definitions. Motif-based haplotype selection supports examination of sequence-defined alleles, whereas coordinate linkage of unmerged paired reads extends the observable CpG span of long amplicons without imputing the intervening sequence. For NGS data, PANDA dereplicates identical reads before alignment and carries their retained counts forward into downstream calculations. Independent sequence variants can be aligned in parallel, although samples containing many unique variants remain computationally demanding. PANDA is therefore intended for targeted amplicon analysis rather than genome-wide mapping.

The single-worker benchmark showed that wall time rose substantially with unique sequence diversity after dereplication: 10,000 distinct patterns required approximately 18 min at 20,000 reads on the tested workstation. Increasing read depth from 10,000 to 50,000 at 3,000 patterns had much less effect on runtime. The tested range does not define a universal recommended read depth or predict performance on other hardware.

Long-read technologies increasingly provide contiguous methylation profiles across extended genomic regions (Flusberg et al., 2010; Simpson et al., 2017; Wenger et al., 2019; Fu et al., 2025; Liu and Conesa, 2025; Montano and Timp, 2025). As high-depth targeted amplicon data remain useful for focused analysis of predefined loci, PANDA complements these approaches by preserving and analyzing read-level methylation patterns from such data.

### Limitations

PANDA analyzes reads from predefined bisulfite amplicons and requires an appropriate reference sequence and consistent preprocessing of the input data. NGS data should undergo adapter and primer trimming, FASTQ quality assessment, and any experiment-specific preprocessing before analysis. Because NGS reads are dereplicated by exact end-to-end sequence identity, inconsistent read boundaries may separate otherwise identical sequences into different variants. Alignment assumes that reads derive from the defined target; motif filtering selects specified post-conversion haplotypes rather than discovering unexpected alleles. Unmerged paired-end analysis requires reliable R1/R2 naming. Automatic strand assignment can be uncertain in heavily contaminated libraries; low-confidence warnings should prompt inspection of the strand diagnostics or explicit reference-orientation selection.

Within-sample heterogeneity estimates depend on coverage, missing CpG calls, read-count weighting, and the number and positions of CpGs shared between records. Changing the minimum shared-CpG threshold affects Amplicon qFDRP only when it changes the set of eligible read pairs; it does not change the CpGs used to calculate the distance for a retained pair. The reported eligibility diagnostics and shared-CpG distribution help identify when threshold sensitivity is relevant. In the unequal-recovery benchmark, count-weighted metrics matched the sampled-read population, whereas unequal molecule recovery shifted that population relative to the original molecules. Giving each distinct variant equal weight did not consistently recover the original-molecule values. As the simulation varied read yield without modeling PCR cycles, it cannot quantify amplification bias in an experimental library. Retained counts should therefore be interpreted as read abundance rather than counts of unique molecules unless independent molecular labels support the latter. Heterogeneity scores from other tools may use different eligibility and weighting rules and should be compared only after these definitions are harmonized. PANDA does not separately report methylation entropy because it summarizes the same four-CpG pattern-frequency distribution used for Window Epipolymorphism, albeit with different sensitivity to rare patterns.

CpG-level tests in PANDA are intended as exploratory comparisons between independent samples. They use sample-level methylation percentages rather than modeling read counts and may be unreliable with few samples or values near 0% or 100%. When evaluating a methylation difference between groups at an individual CpG, the *p*-value should be considered alongside the sample-level percentages, the magnitude of the difference, and the experimental design.

Coordinate linkage of unmerged paired reads preserves the CpG calls observed at both ends of an amplicon, but methylation states in the unsequenced interval cannot be inferred. Long-read sequencing can provide continuous methylation profiles when coverage across a larger region is needed (Flusberg et al., 2010; Simpson et al., 2017; Wenger et al., 2019).

## Conclusion

The synthetic demonstrations, controlled benchmarks, and public-data re-analysis characterize PANDA as a framework for phased, read-level analysis of targeted bisulfite amplicons. By preserving physical CpG coordinates and observed NGS read abundance, it allows methylation-pattern structure and within-sample heterogeneity to be evaluated alongside conventional CpG-level estimates, with interpretation guided by filtering, read span, and experimental design.

## Statements

### Data availability statement

The PANDA source code, graphical application, command line workflow, PANDAcore package, documentation, and synthetic demonstration materials are available at https://github.com/kubo-azu/PANDA under the MIT License. Versioned releases and the dependency environment recorded in `renv.lock` support reproducible local execution, and the hosted graphical interface is available at https://huggingface.co/spaces/kubo-azu/PANDA. The *ST8SIA1* TBAS data used in the re-analysis were obtained from the supplementary material accompanying the source publication (Bogutz et al., 2019).

### Ethics statement

Ethical review and approval were not required because this study used synthetic datasets and re-analyzed publicly available data.

### Author contributions

AK: Conceptualization, Data curation, Formal analysis, Funding acquisition, Investigation, Software, Validation, and Writing – original draft. HK: Conceptualization, Investigation, Resources, Supervision, Validation, and Writing – review and editing. AT: Resources, Supervision, Validation, and Writing – review and editing.

## Acknowledgments

The authors thank Drs. Kazuhiko Nakabayashi, Shin-ichi Horike, and Shiori Minabe for their constructive feedback, which contributed to the development of PANDA.

## Generative AI statement

The author(s) declared that generative AI was not used in the creation of this manuscript.

## Supplementary material

**Figure S1.** Global methylation estimates in synthetic datasets. (Fig_S1.pdf)

**Figure S2.** Heterogeneity metrics in the synthetic Sanger datasets. (Fig_S2.pdf)

**Figure S3.** Representative Sanger methylation patterns in synthetic Experiment A. (Fig_S3.pdf)

**Figure S4.** Differential read recovery and weighting of heterogeneity estimates. (Fig_S4.pdf)

**Figure S5.** Computational scaling with distinct input sequences and read depth. (Fig_S5.pdf)

**Figure S6.** Abundance heatmaps for the additional *ST8SIA1* locus samples. (Fig_S6.pdf)

**Table S1.** PANDA software environment and dependencies. (Table_S1.xlsx)

**Table S2.** Effects of conversion and substitution errors in controlled *H19* benchmarks. (Table_S2.xlsx)

**Table S3.** Effects of motif length and matching rule on haplotype assignment in controlled *H19* benchmarks. (Table_S3.xlsx)

**Table S4.** Summary of the public *ST8SIA1* locus re-analysis results. (Table_S4.xlsx)

**Data S1.** Controlled synthetic benchmark datasets and generation scripts. (Data_S1.zip)

**Data S2.** Synthetic demonstration design and per-sample results. (Data_S2.xlsx)

**Data S3.** Numerical results underlying the synthetic-data validation and CpG-level group comparisons. (Data_S3.xlsx)

## References

Adusumalli, S., Mohd Omar, M. F., Soong, R., and Benoukraf, T. (2015). Methodological aspects of whole-genome bisulfite sequencing analysis. Brief. Bioinform. 16, 369–379. doi: 10.1093/bib/bbu016.

Bock, C., Reither, S., Mikeska, T., Paulsen, M., Walter, J., and Lengauer, T. (2005). BiQ Analyzer: visualization and quality control for DNA methylation data from bisulfite sequencing. Bioinformatics 21, 4067–4068. doi: 10.1093/bioinformatics/bti652.

Bogutz, A. B., Brind’Amour, J., Kobayashi, H., Jensen, K. N., Nakabayashi, K., Imai, H., et al. (2019). Evolution of imprinting via lineage-specific insertion of retroviral promoters. Nat. Commun. 10, 5674. doi: 10.1038/s41467-019-13662-9.

Bolger, A. M., Lohse, M., and Usadel, B. (2014). Trimmomatic: a flexible trimmer for Illumina sequence data. Bioinformatics 30, 2114–2120. doi: 10.1093/bioinformatics/btu170.

Chen, S., Zhou, Y., Chen, Y., and Gu, J. (2018). fastp: an ultra-fast all-in-one FASTQ preprocessor. Bioinformatics 34, i884–i890. doi: 10.1093/bioinformatics/bty560.

Chen, S. (2023). Ultrafast one-pass FASTQ data preprocessing, quality control, and deduplication using fastp. iMeta 2, e107. doi: 10.1002/imt2.107.

Farrell, C., Thompson, M., Tosevska, A., Oyetunde, A., and Pellegrini, M. (2021). BiSulfite Bolt: A bisulfite sequencing analysis platform. GigaScience 10, giab033. doi: 10.1093/gigascience/giab033.

Flusberg, B. A., Webster, D. R., Lee, J. H., Travers, K. J., Olivares, E. C., Clark, T. A., et al. (2010). Direct detection of DNA methylation during single-molecule, real-time sequencing. Nat. Methods 7, 461–465. doi: 10.1038/nmeth.1459.

Fu, Y., Timp, W., and Sedlazeck, F. J. (2025). Computational analysis of DNA methylation from long-read sequencing. Nat. Rev. Genet. 26, 620–634. doi: 10.1038/s41576-025-00822-5.

Guo, W., Fiziev, P., Yan, W., Cokus, S., Sun, X., Zhang, M. Q., et al. (2013). BS-Seeker2: a versatile aligning pipeline for bisulfite sequencing data. BMC Genomics 14, 774. doi: 10.1186/1471-2164-14-774.

Jenkinson, G., Pujadas, E., Goutsias, J., and Feinberg, A. P. (2017). Potential energy landscapes identify the information-theoretic nature of the epigenome. Nat. Genet. 49, 719–729. doi: 10.1038/ng.3811.

Krueger, F., and Andrews, S. R. (2011). Bismark: a flexible aligner and methylation caller for Bisulfite-Seq applications. Bioinformatics 27, 1571–1572. doi: 10.1093/bioinformatics/btr167.

Kumaki, Y., Oda, M., and Okano, M. (2008). QUMA: quantification tool for methylation analysis. Nucleic Acids Res. 36, W170–W175. doi: 10.1093/nar/gkn294.

Laisné, M., Lupien, M., and Vallot, C. (2025). Epigenomic heterogeneity as a source of tumour evolution. Nat. Rev. Cancer 25, 7–26. doi: 10.1038/s41568-024-00757-9.

Landan, G., Cohen, N. M., Mukamel, Z., Bar, A., Molchadsky, A., Brosh, R., et al. (2012). Epigenetic polymorphism and the stochastic formation of differentially methylated regions in normal and cancerous tissues. Nat. Genet. 44, 1207–1214. doi: 10.1038/ng.2442.

Landau, D. A., Clement, K., Ziller, M. J., Boyle, P., Fan, J., Gu, H., et al. (2014). Locally disordered methylation forms the basis of intratumor methylome variation in chronic lymphocytic leukemia. Cancer Cell 26, 813–825. doi: 10.1016/j.ccell.2014.10.012.

Lewin, J., Schmitt, A. O., Adorján, P., Hildmann, T., and Piepenbrock, C. (2004). Quantitative DNA methylation analysis based on four-dye trace data from direct sequencing of PCR amplificates. Bioinformatics 20, 3005–3012. doi: 10.1093/bioinformatics/bth346.

Liu, T., and Conesa, A. (2025). Profiling the epigenome using long-read sequencing. Nat. Genet. 57, 27–41. doi: 10.1038/s41588-024-02038-5.

Lutsik, P., Feuerbach, L., Arand, J., Lengauer, T., Walter, J., and Bock, C. (2011). BiQ Analyzer HT: locus-specific analysis of DNA methylation by high-throughput bisulfite sequencing. Nucleic Acids Res. 39, W551–W556. doi: 10.1093/nar/gkr312.

Masser, D. R., Berg, A. S., and Freeman, W. M. (2013). Focused, high accuracy 5-methylcytosine quantitation with base resolution by benchtop next-generation sequencing. Epigenetics Chromatin 6, 33. doi: 10.1186/1756-8935-6-33.

Mattei, A. L., Bailly, N., and Meissner, A. (2022). DNA methylation: a historical perspective. Trends Genet. 38, 676–707. doi: 10.1016/j.tig.2022.03.010.

Meaburn, E. L., Schalkwyk, L. C., and Mill, J. (2010). Allele-specific methylation in the human genome: implications for genetic studies of complex disease. Epigenetics 5, 578–582. doi: 10.4161/epi.5.7.12960.

Meissner, A., Gnirke, A., Bell, G. W., Ramsahoye, B., Lander, E. S., and Jaenisch, R. (2005). Reduced representation bisulfite sequencing for comparative high-resolution DNA methylation analysis. Nucleic Acids Res. 33, 5868–5877. doi: 10.1093/nar/gki901.

Montano, C., and Timp, W. (2025). Evolution of genome-wide methylation profiling technologies. Genome Res. 35, 572–582. doi: 10.1101/gr.278407.123.

Needleman, S. B., and Wunsch, C. D. (1970). A general method applicable to the search for similarities in the amino acid sequence of two proteins. J. Mol. Biol. 48, 443–453. doi: 10.1016/0022-2836(70)90057-4.

Nikolaienko, O., Lønning, P. E., and Knappskog, S. (2023). epialleleR: an R/Bioconductor package for sensitive allele-specific methylation analysis in NGS data. GigaScience 12, giad087. doi: 10.1093/gigascience/giad087.

Rosenski, J., Peretz, A., Magenheim, J., Loyfer, N., Shemer, R., Glaser, B., et al. (2025). Atlas of imprinted and allele-specific DNA methylation in the human body. Nat. Commun. 16, 2141. doi: 10.1038/s41467-025-57433-1.

Scala, G., Affinito, O., Palumbo, D., Florio, E., Monticelli, A., Miele, G., et al. (2016). ampliMethProfiler: a pipeline for the analysis of CpG methylation profiles of targeted deep bisulfite sequenced amplicons. BMC Bioinformatics 17, 484. doi: 10.1186/s12859-016-1380-3.

Scherer, M., Nebel, A., Franke, A., Walter, J., Lengauer, T., Bock, C., et al. (2020). Quantitative comparison of within-sample heterogeneity scores for DNA methylation data. Nucleic Acids Res. 48, e46. doi: 10.1093/nar/gkaa120.

Simpson, J. T., Workman, R. E., Zuzarte, P. C., David, M., Dursi, L. J., and Timp, W. (2017). Detecting DNA cytosine methylation using nanopore sequencing. Nat. Methods 14, 407–410. doi: 10.1038/nmeth.4184.

Smith, Z. D., Hetzel, S., and Meissner, A. (2025). DNA methylation in mammalian development and disease. Nat. Rev. Genet. 26, 7–30. doi: 10.1038/s41576-024-00760-8.

Tycko, B. (2010). Allele-specific DNA methylation: beyond imprinting. Hum. Mol. Genet. 19, R210–R220. doi: 10.1093/hmg/ddq376.

Vaisvila, R., Ponnaluri, V. K. C., Sun, Z., Langhorst, B. W., Saleh, L., Guan, S., et al. (2021). Enzymatic methyl sequencing detects DNA methylation at single-base resolution from picograms of DNA. Genome Res. 31, 1280–1289. doi: 10.1101/gr.266551.120.

Wang, K., Liu, H., Hu, Q., Wang, L., Liu, J., Zheng, Z., et al. (2022). Epigenetic regulation of aging: implications for interventions of aging and diseases. Signal Transduct. Target. Ther. 7, 374. doi: 10.1038/s41392-022-01211-8.

Wenger, A. M., Peluso, P., Rowell, W. J., Chang, P.-C., Hall, R. J., Concepcion, G. T., et al. (2019). Accurate circular consensus long-read sequencing improves variant detection and assembly of a human genome. Nat. Biotechnol. 37, 1155–1162. doi: 10.1038/s41587-019-0217-9.

Wong, N. C., Pope, B. J., Candiloro, I. L., Korbie, D., Trau, M., Wong, S. Q., et al. (2016). MethPat: a tool for the analysis and visualisation of complex methylation patterns obtained by massively parallel sequencing. BMC Bioinformatics 17, 98. doi: 10.1186/s12859-016-0950-8.

Yücel, A. D., and Gladyshev, V. N. (2026). Systemic epigenetic dysregulation as a driver of ageing and a therapeutic target. Nat. Rev. Mol. Cell Biol. 27, 528–542. doi: 10.1038/s41580-026-00958-0.

Zoghbi, H. Y., and Beaudet, A. L. (2016). Epigenetics and Human Disease. Cold Spring Harb. Perspect. Biol. 8, a019497. doi: 10.1101/cshperspect.a019497.

