## Supplementary material for "PANDA: Phased Analysis of DNA Amplicons for Read-Level Methylation Profiling and Heterogeneity Assessment": Fig_S1

**A**

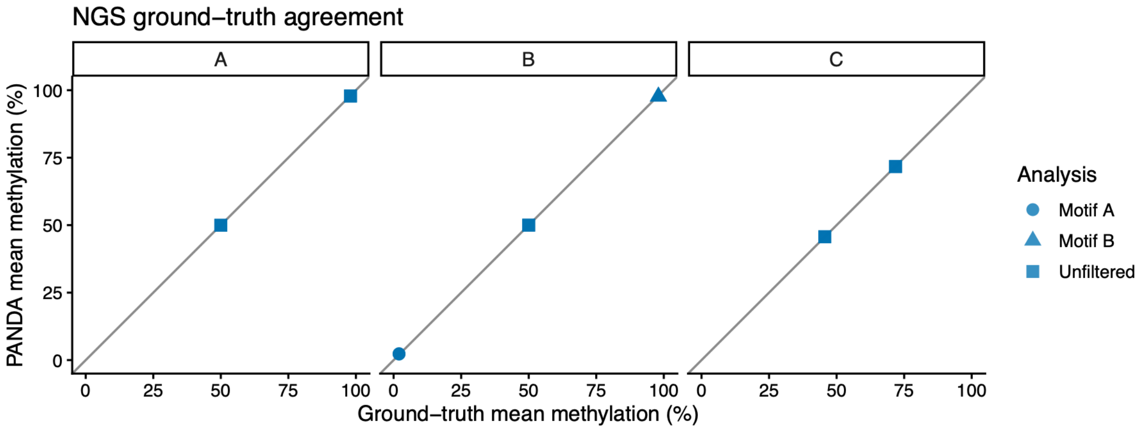


**B**

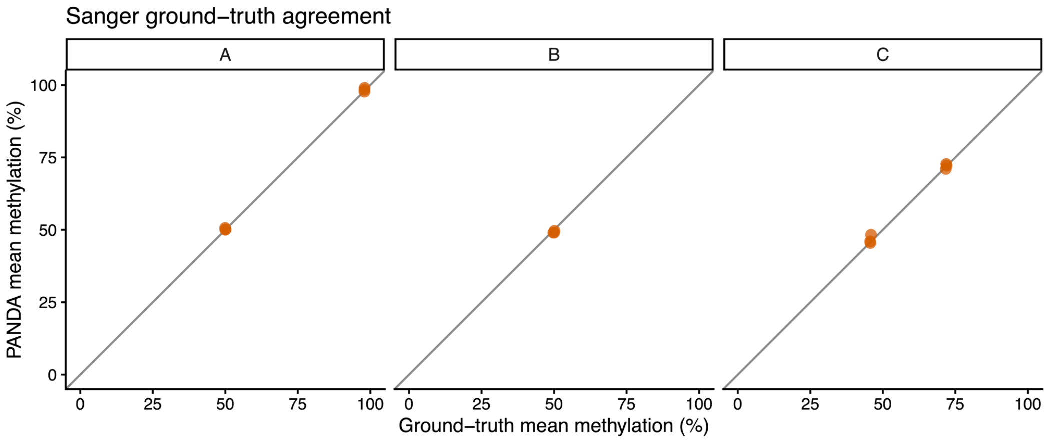


**Figure S1. Agreement between predefined and PANDA-estimated global methylation in the synthetic datasets.** Ground-truth mean methylation is plotted against the corresponding PANDA estimate for the amplicon-NGS **(A)** and Sanger **(B)** datasets. Within each plot, facets A–C correspond to synthetic Experiments A–C, respectively. The NGS plot additionally distinguishes unfiltered and motif-selected analyses in Experiment B. Each point represents an independently generated replicate, and the diagonal line indicates exact agreement.
