## Supplementary material for "PANDA: Phased Analysis of DNA Amplicons for Read-Level Methylation Profiling and Heterogeneity Assessment": Fig_S2

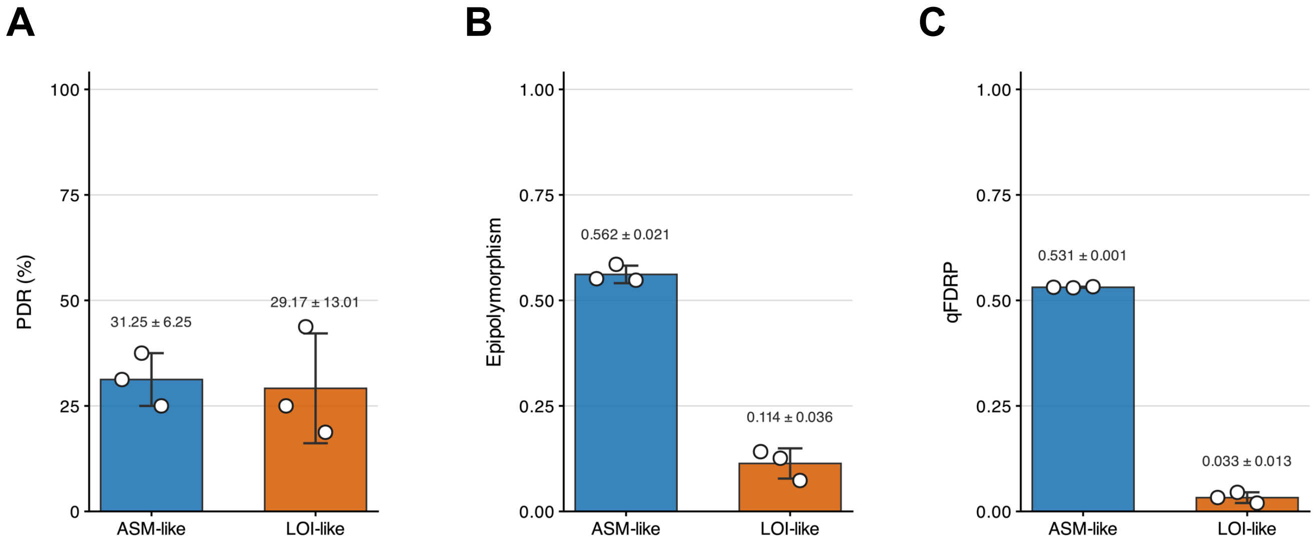


**Figure S2. Heterogeneity metrics in the synthetic Sanger datasets.**

Replicate-level Amplicon PDR **(A)**, Window Epipolymorphism **(B)**, and Amplicon qFDRP **(C)** are shown across the predefined synthetic methylation states. Each Sanger clone contributes equally with Count = 1.
