## Supplementary material for "PANDA: Phased Analysis of DNA Amplicons for Read-Level Methylation Profiling and Heterogeneity Assessment": Fig_S3

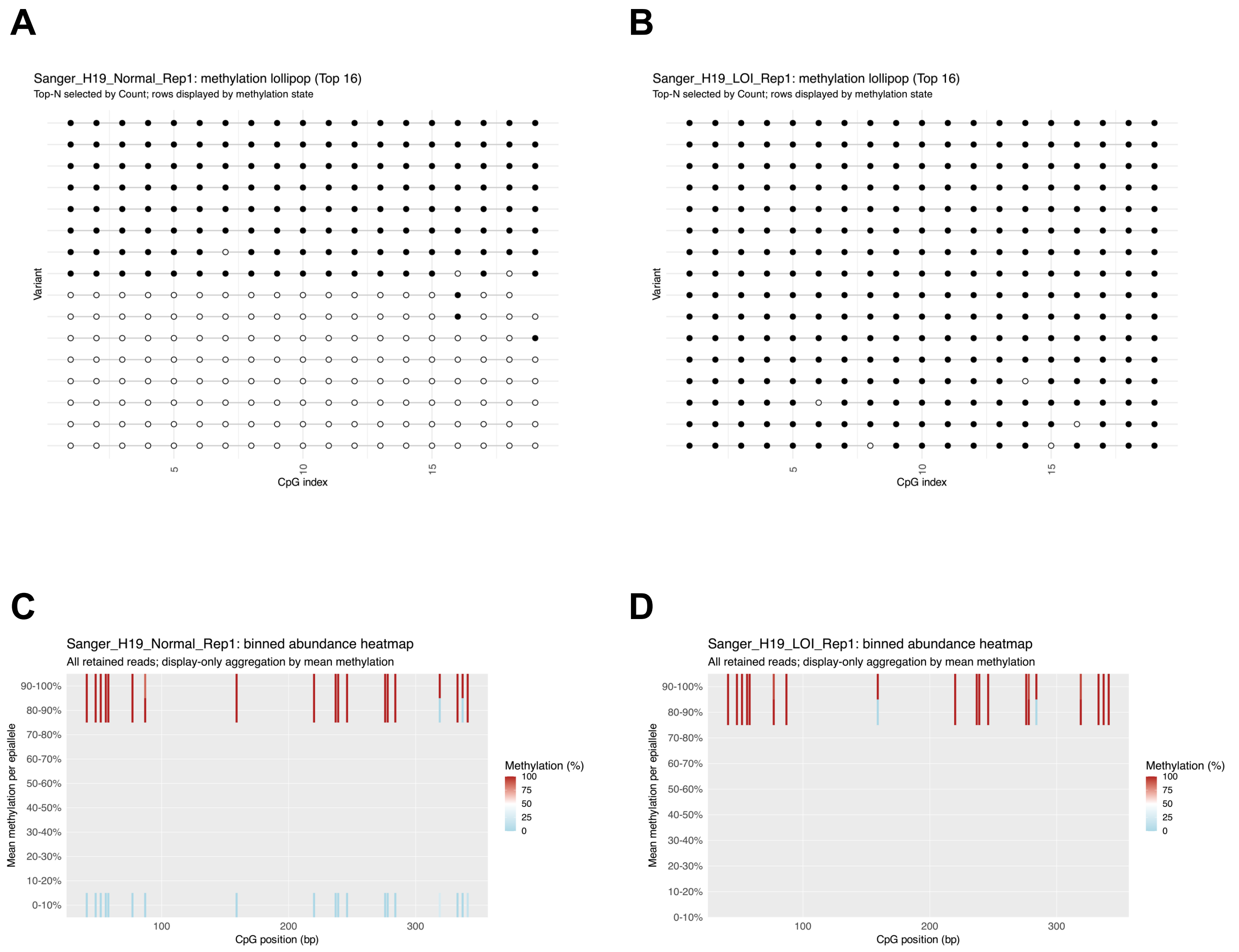


**Figure S3. Representative Sanger methylation patterns in synthetic Experiment A.**

Ranked clone-level lollipop plots are shown for representative ASM-like *H19* Normal **(A)** and LOI-like *H19* **(B)** samples. Corresponding binned abundance heatmaps are shown in **(C)** and **(D)**, respectively. Each Sanger clone contributes equally with Count = 1.
