## Supplementary material for "PANDA: Phased Analysis of DNA Amplicons for Read-Level Methylation Profiling and Heterogeneity Assessment": Fig_S4

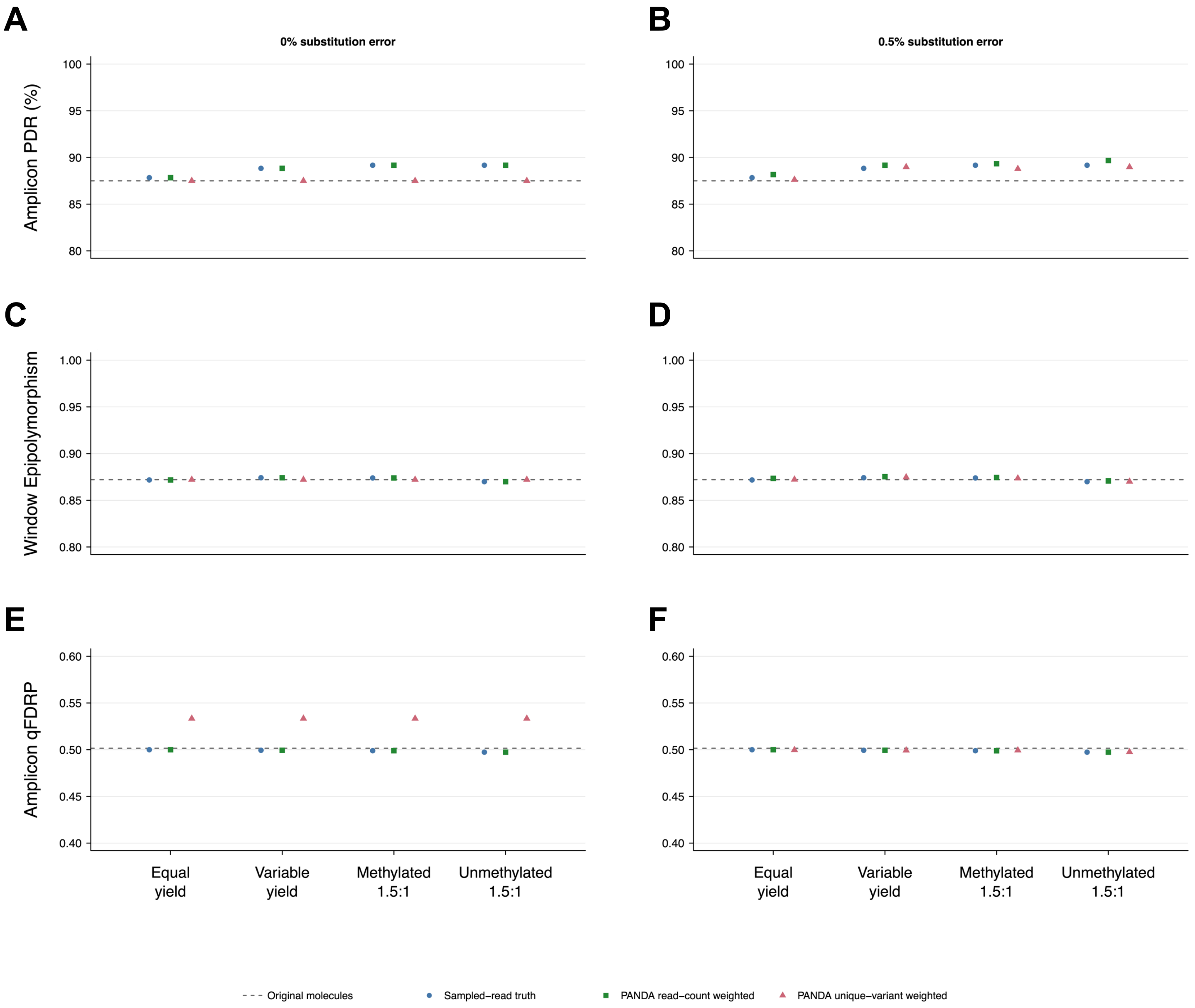


**Figure S4. Differential read recovery and weighting of heterogeneity estimates.**

**(A,C,E)** No substitution error; **(B,D,F)** 0.5% per-base substitution error. Rows show Amplicon PDR, Window Epipolymorphism, and Amplicon qFDRP. Six hundred reads were sampled from 320 *H19*-based molecules under equal yield, Gamma-distributed variable yield, or preferential recovery of methylated or unmethylated molecules. In the latter conditions, fully methylated versus fully unmethylated molecules had relative selection weights of 1.5:1 or 1:1.5. The dashed line shows the original-molecule value; circles show sampled-read truth; squares show PANDA estimates weighted by retained read count; triangles show estimates after assigning equal weight to each distinct retained sequence variant.
