## Supplementary material for "PANDA: Phased Analysis of DNA Amplicons for Read-Level Methylation Profiling and Heterogeneity Assessment": Fig_S5

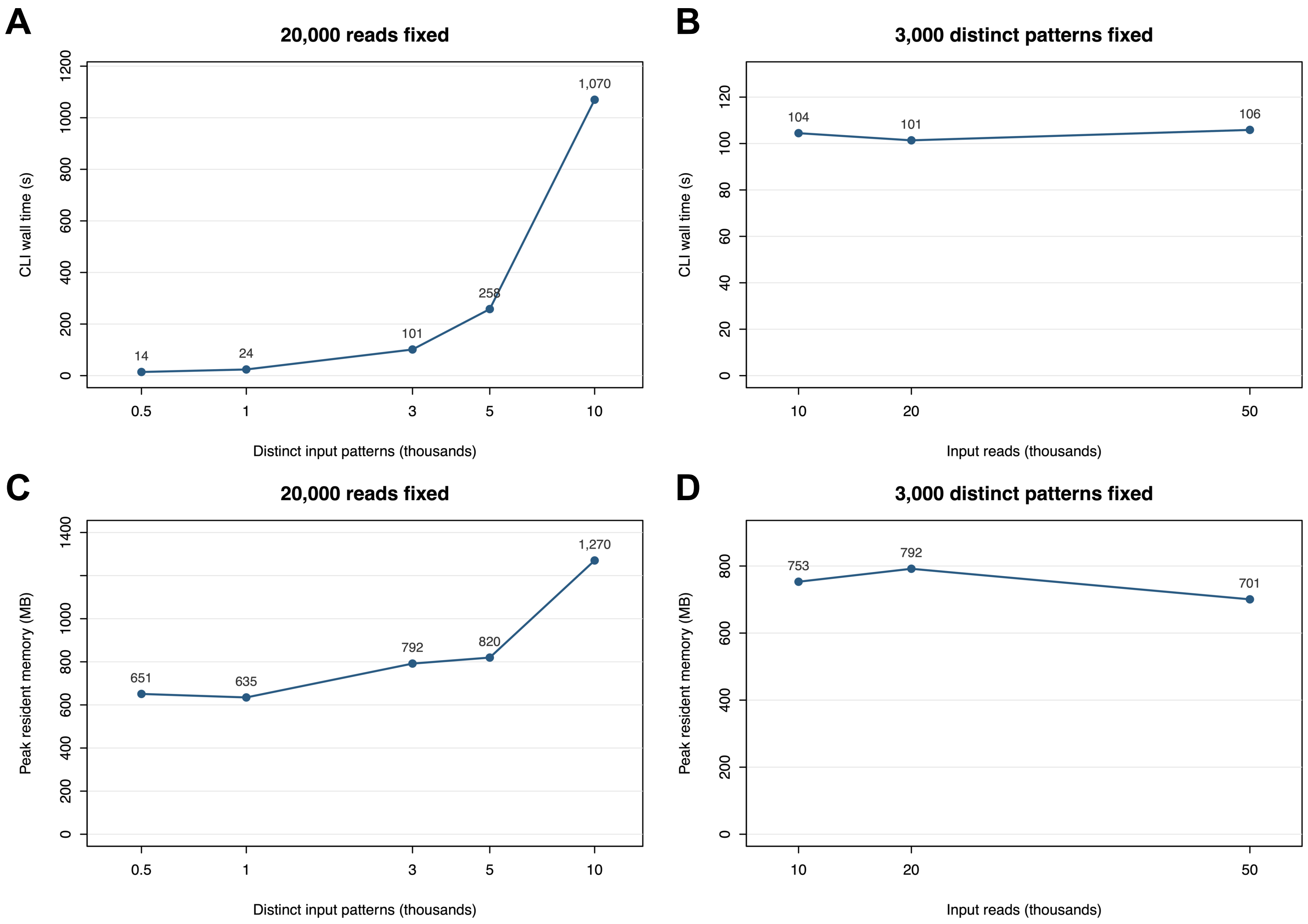


**Figure S5. Computational scaling with distinct input sequences and read depth.**

**(A,C)** Twenty thousand reads with 500–10,000 distinct input sequences; the horizontal axes are logarithmic. **(B,D)** Three thousand distinct input sequences with 10,000–50,000 reads. Panels A and B show single-worker PANDA CLI wall time, including R startup; panels C and D show peak resident memory. Each point represents one run on the hardware specified in Methods; lines connect the measured conditions.
