## Supplementary material for "PANDA: Phased Analysis of DNA Amplicons for Read-Level Methylation Profiling and Heterogeneity Assessment": Fig_S6

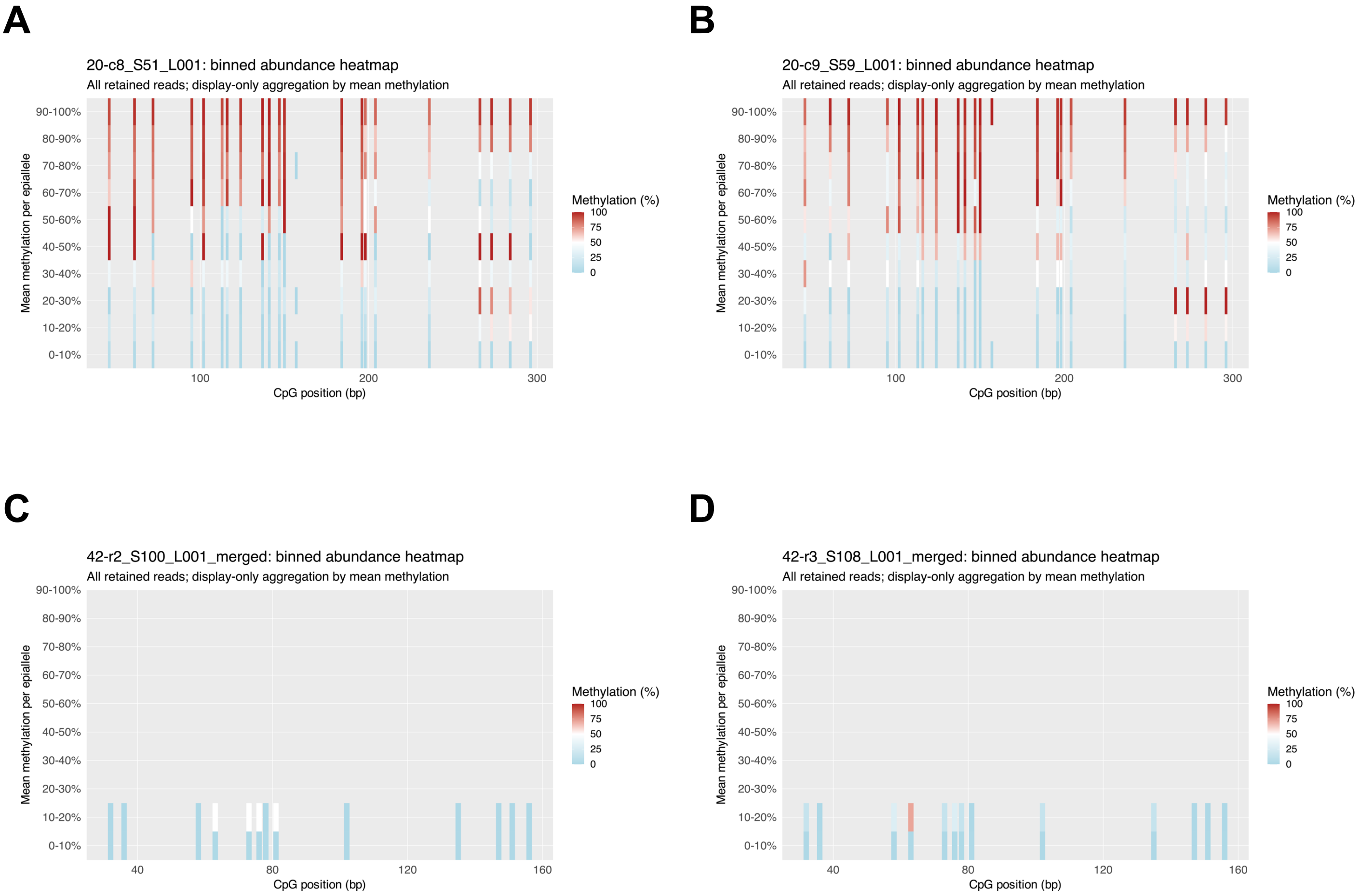


**Figure S6. Abundance heatmaps for the additional *ST8SIA1* locus samples.**

Binned abundance heatmaps are shown for two additional coordinate-linked chimpanzee unmerged R1/R2 samples **(A,B)** and two additional fastp-merged rhesus macaque samples **(C,D)**. Each heatmap was generated from all retained reads using read-count weights and physical CpG coordinates. The representative samples are shown in Figure 7.
